# Interplay between ß-propeller subunits WDR26 and muskelin regulates the CTLH E3 ligase supramolecular complex

**DOI:** 10.1101/2024.03.08.584134

**Authors:** Matthew E.R. Maitland, Gabriel Onea, Dominic D. G. Owens, Brianna C. Gonga-Cavé, Xu Wang, Cheryl H. Arrowsmith, Dalia Barsyte-Lovejoy, Gilles A. Lajoie, Caroline Schild-Poulter

## Abstract

The Pro/N-degron recognizing C-terminal to LisH (CTLH) complex is an E3 ligase of emerging interest in the developmental field and for targeted protein degradation (TPD) modalities. The human CTLH complex forms distinct supramolecular ring-shaped structures dependent on the multimerization of WDR26 or muskelin ß-propeller proteins. Here, we find that, in human cells, CTLH complex E3 ligase activity is dictated by a dynamic exchange between WDR26 and muskelin in tandem with muskelin autoregulation. Proteomic experiments revealed that complex-associated muskelin protein turnover is a major ubiquitin-mediated degradation event dependent on the CTLH complex in unstimulated HeLa cells. We observed that muskelin and WDR26 binding to the scaffold of the complex is interchangeable, indicative of the formation of separate WDR26 and muskelin complexes, which correlated with distinct proteomes in WDR26 and muskelin knockout cells. We found that mTOR inhibition-induced degradation of Pro/N-degron containing protein HMGCS1 is distinctly regulated by a muskelin-specific CTLH complex. Finally, we found that mTOR inhibition also activated muskelin degradation, likely as an autoregulatory feedback mechanism to regulate CTLH complex activity. Thus, rather than swapping substrate receptors, the CTLH E3 ligase complex controls substrate selectivity and its autoregulation through exchanging its β-propeller oligomeric subunits WDR26 and muskelin.

## Introduction

Ubiquitination involves the covalent attachment of one or more ubiquitin (Ub) proteins to mainly lysine residues of substrate proteins. Depending on the type of ubiquitin linkage and topology of the poly-Ub chain, various effects on the targeted substrate can occur such as proteasomal degradation, changes in interaction partners, or alterations of subcellular localization (Komander and Rape 2012; Dikic and Schulman 2023). These changes to substrate function and signalling pathways are crucial for proper metazoan development; for this reason, a dysfunctional ubiquitination cascade leads to aberrant cellular differentiation and developmental abnormalities, among other pathologies (Rape 2017; Popovic et al. 2014).

The transfer of Ub to a substrate is achieved through the coordinated function of E1 Ub-activating enzymes, E2 Ub-conjugating enzymes, and E3 Ub-protein ligases (Pickart 2001). The majority of E3 ligases contain RING (Really Interesting New Gene) domains that confer substrate specificity by bridging the interaction between the E2∼Ub conjugate and a specific substrate. Human cells potentially express over 600 predicted RING E3 ligases to regulate a variety of molecular pathways - many of which exist in multi-subunit complexes comprising interchangeable substrate receptors, adaptors, scaffolds, and RING catalytic modules - thereby increasing the theoretical number of proteins that could be regulated through ubiquitination (Deshaies and Joazeiro 2009; Vittal et al. 2015)

The C-terminal to LisH (CTLH) complex – also known as the Glucose-Induced Degradation Deficient (GID) complex – is a multi-subunit RING E3 ligase that has been implicated in a growing list of biological processes such as cell proliferation, haemopoietic differentiation, glycolysis, autophagy, and maternal to zygotic transition (Maitland et al. 2022; McTavish et al. 2019; Maitland et al. 2021; Lampert et al. 2018; Liu et al. 2020; Sherpa et al. 2022; Zavortink et al. 2020; Cao et al. 2020; Pfirrmann et al. 2015; Santt et al. 2008). Recent studies have elucidated the composition and general structure of the CTLH complex *in vitro* using microscopy and *in vitro* binding assays (van gen Hassend et al. 2023; Qiao et al. 2020; Sherpa et al. 2021; Mohamed et al. 2021). Briefly, E3 ligase activity is mediated by the RING catalytic module formed between a MAEA/RMND5A heterodimer where each monomer binds to a separate GID8 (TWA1) protein. The general scaffold is comprised of GID8, RanBP9 (RanBPM), and ARMC8 isoforms α and β. The Pro/N-degron recognizing substrate receptor GID4 then binds to ARMC8α, resulting in a horseshoe-shaped structure with the RMND5A-MAEA heterodimer linking two separate CTLH scaffold + GID4 moieties, with one RanBP9 protein at each free end. Addition of WDR26 promotes the formation of a 1.5 MDa supramolecular donut-shaped complex by forming 2 homodimers and connecting the 2 horseshoe-shaped structures through RanBP9 on opposite ends of the complex. This supramolecular complex can accommodate substrates in its internal cavity, leading to their ubiquitination (Sherpa et al. 2021; Qiao et al. 2020).

Interestingly, certain species, including mammals, have another CTLH complex member called muskelin (Maitland et al. 2022, 2019). Similar to WDR26, this protein contains a β-propeller-like Kelch domain at its C-terminus. *In vitro*, muskelin was also shown to mediate the oligomerization of the CTLH complex through a homotetramer instead of a homodimer, potentially creating a different size central pocket for oligomeric substrates (Sherpa et al. 2021; van gen Hassend et al. 2023).

Although *in vitro* muskelin and WDR26 can both mediate the formation of CTLH complex oligomerization, in cells, only the loss of WDR26, and not muskelin, has a profound effect on CTLH complex formation (Sherpa et al. 2021; Onea et al. 2022). An additional distinction is that muskelin protein levels, but not WDR26, are regulated by ubiquitination in a MAEA/RMND5A-dependent manner (Maitland et al. 2019; Jordan et al. 2023). Despite differences between these two structurally-similar subunits, it remains unclear whether they translate into differences in CTLH complex-mediated regulation. In this study, using proteomics and complex formation assays in human cells, we found that WDR26 and muskelin can mediate the formation of two distinct supramolecular CTLH complexes that have distinct functions. We also discovered that inhibition of mTOR signalling regulates muskelin-specific CTLH complex activity in human cells and leads to muskelin autoregulation.

## Results

### Muskelin autoregulation is a dominant activity of the CTLH complex

To examine possible ubiquitination targets of the CTLH complex, we conducted global label-free quantitative proteomics in RMND5A (n=5) or MAEA (n=4) HeLa knockout cells. We reasoned that degradation targets of the CTLH complex would be increased in RMND5A or MAEA KO cells relative to control cells. Principal components analysis and hierarchical clustering revealed that knockouts clustered distinctly from control samples (**Supplemental Figure 1A**). Out of 4769 quantified proteins, 202 were significantly increased in RMND5A KO and 181 proteins were increased in MAEA KO samples compared to controls (adjusted p<0.05, KO/WT fold change>1.5) (**Figure 1A, B, Supplemental data 1**). Overall, there was substantial agreement between changed proteins in both knockouts, with 108 significantly changed proteins identified in common (**Figure 1C**).

**Figure 1.**
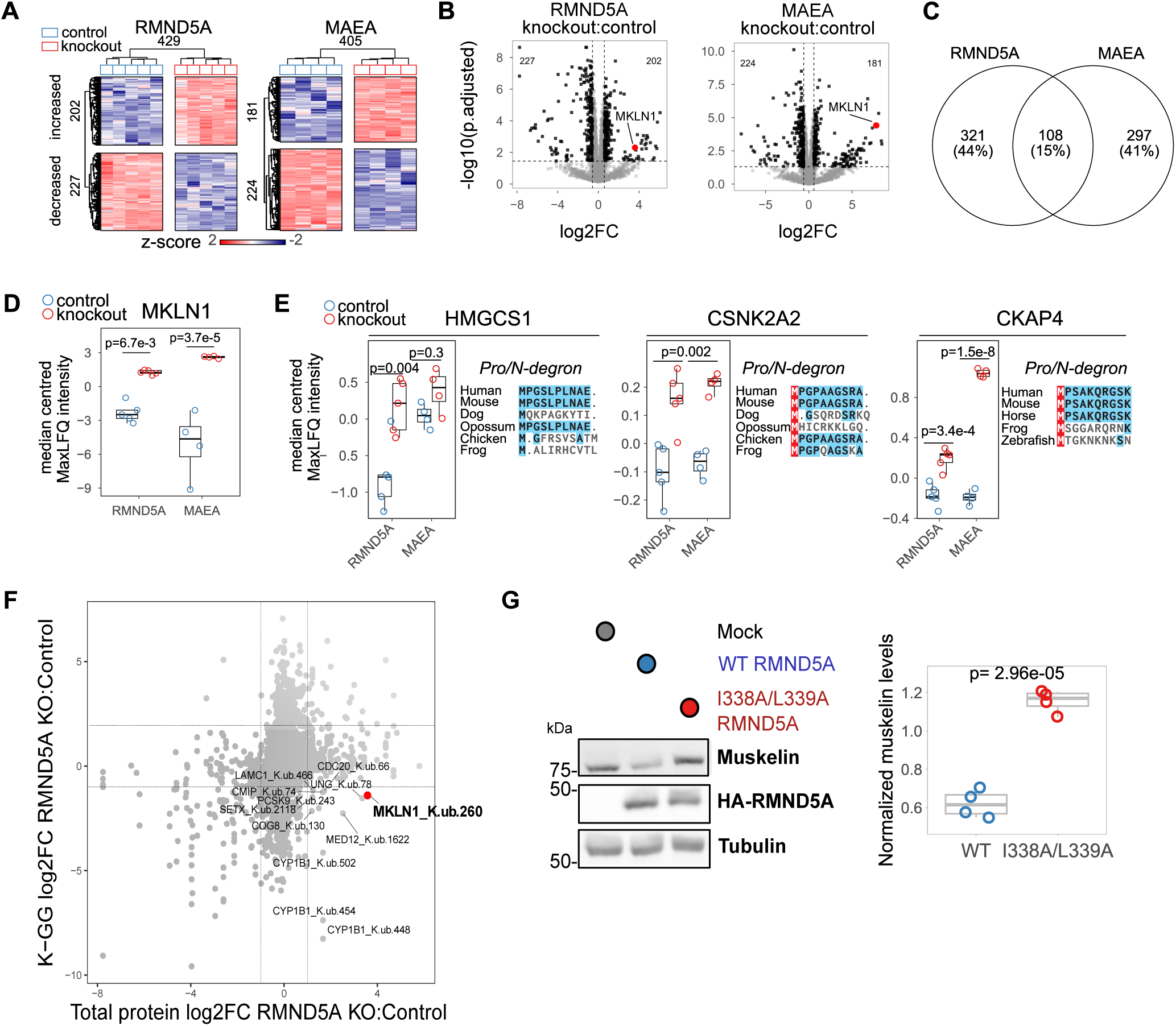
Muskelin is the primary ubiquitination target of the CTLH complex in unstimulated HeLa cells. A) Heatmap of differentially abundant proteins (adjusted p-value < 0.05, fold change > 1.5, statistics derived from limma, see methods) in whole-cell extracts of RMND5A (n=5) or MAEA KO (n=4) HeLa cells. Each row represents one protein and samples are represented in columns. Protein abundance is shown as z-scores quantified across rows. Hierarchical clustering of samples and proteins is shown. B) Volcano plot of all proteins with log2 fold change shown on the x-axis and −log10 adjusted p values shown on the y axis. RMND5A KO vs respective controls on the left, MAEA KO vs respective controls on the right. Muskelin (MKLN1) is indicated in red. C) Overlap of proteins that had significant differential abundance in RMND5A or MAEA KO cells. D) Median centered intensity of muskelin (MKLN1) in the proteome samples. Boxplot midline indicates median values, bounds of the box indicate 25th and 75th percentiles, and maxima and minima indicate the largest point above or below 1.5 times the interquartile range. E) Median centered intensities for specific proteins containing Pro/N-degrons known to by recognized by GID4. Evolutionary conservation of first ten amino acids of N-termini are indicated. Boxplot midline indicates median values, bounds of the box indicate 25th and 75th percentiles, and maxima and minima indicate the largest point above or below 1.5 times the interquartile range. F) Comparison of the diGLY-enriched proteome vs total proteome in RMND5A KO HeLa cells. MG-132-treated control or RMND5A KO HeLa cells were lysed, digested, and peptides containing the diGLY motif originating from ubiquitin were enriched and analyzed by LC-MS/MS. Shown are diGLY peptides plotted with their log2 fold change on the y-axis against the log2 fold change of their matched total protein levels. diGLY peptides with log2FC>1 are labelled with protein name and modified Lysine residue position. Muskelin K260 is indicated in red. G) Restoration of muskelin levels in RMND5A KO cells by exogenous WT RMND5A or I338A/L399A E2 binding deficient mutant. RMND5A KO HeLa cells were transfected with carrier plasmid (mock), HA-WT-RMND5A, or HA-I338A/L339A-RMND5A. Whole cell lysates were prepared 24 hours later and analyzed by western blot with the indicated antibodies. Boxplot midline indicates median values of muskelin intensity (normalized to tubulin, relative to mock for each replicate), bounds of the box indicate 25th and 75th percentiles, and maxima and minima indicate the largest point above or below 1.5 times the interquartile range (n=4). Welch two sample t-test, t = 11.611, df ≈ 6.

We previously discovered that protein levels of CTLH complex subunit muskelin (MKLN1) are increased in RMND5A and MAEA KO HeLa and HEK293 cell lines (Maitland et al. 2019). Interestingly, muskelin was one of the most highly increased proteins of the quantifiable proteome for both RMND5A and MAEA KO cells (**Figure 1B, D**). Additionally, alanyl aminopeptidase (ANPEP), BAR/IMD domain containing adaptor protein 2 (BAIAP2), and LIMA1 (LIM domain and actin binding 1), all previously identified as interactors of CTLH complex subunits (Onea et al. 2022; Huttlin et al. 2021), were significantly increased in one or both of the knockout cell lines (**Supplemental Figure 1B**). We also noted that some proteins increased in RMND5A and/or MAEA knockouts contain an N-terminal sequence that matches the Pro/N-degron recognized by CTLH complex substrate receptor GID4 (**Figure 1E**) (Dong et al. 2018; Chrustowicz et al. 2022; Chen et al. 2017). This included mevalonate pathway enzyme HMG-CoA synthase 1 (HMGCS1), which was recently identified as a GID4 substrate (Owens & Maitland et al., 2023), and two other proteins not previously linked to GID4 or the CTLH complex: CSNK2A2, which encodes the α’ subunit of casein kinase II (CK2), and cytoskeleton-associated protein 4 (CKAP4), which is associated with cell migration and metastasis (Sun et al. 2022; Osugi et al. 2019).

A comparison of the RMND5A-dependent proteome and ubiquitinome showed few changes consistent with degradative ubiquitination (**Figure 1F, Supplemental data 2, Supplemental Figure 1C, D**), which is in line with our previous observations for CTLH complex scaffolding subunit RanBP9 (Maitland et al. 2021). In fact, muskelin was the top changed protein increased at the protein level with a more than 2-fold decreased ubiquitin site (**Figure 1F**). This occurred on lysine 260 of muskelin, a conserved and – based on the Alphafold predicted structure – surface exposed residue located in the connecting region between the CTLH motif and the kelch repeats (**Supplemental Figure 1E, F)**. This suggests that K260 is at least one site of RMDN5A-dependent ubiquitination of muskelin. Next, we used RMND5A mutants to establish that the muskelin regulation is mediated by the E3 ligase activity of the complex. We assessed muskelin protein levels in RMND5A KO cells, where it is increased, and we compared the effect of transfecting wildtype RMND5A with a I338A/L339A mutant previously shown to be deficient in E2 binding *in vitro* (Sherpa et al. 2021). While exogenous WT RMND5A reduced muskelin protein levels, the I338A/L339A mutant had no effect, thus establishing that the regulation of muskelin is due to the ubiquitin ligase activity of the CTLH complex (**Figure 1G**). Together with our previous studies on global CTLH functions (Maitland et al. 2021; Onea et al. 2022; Owens et al. 2023), these data suggest that in unstimulated HeLa cells, most of the CTLH-dependent ubiquitination is non-degradative, but of the primary degradative ubiquitination target is its own subunit muskelin.

### Muskelin and WDR26 both engage in similar interactions with RanBP9

Intrigued by why the CTLH complex E3 activity would be mainly dedicated to regulating muskelin levels in unstimulated conditions, but not those of WDR26, we sought to compare functions and interactions of WDR26 and muskelin. Studies using *in vitro* reconstituted systems have determined that, in the presence of muskelin or WDR26, both of which bind RanBP9, a supramolecular CTLH complex is formed (Sherpa et al. 2021; van gen Hassend et al. 2023; Mohamed et al. 2021). To establish the requirements for this in a cellular model, we evaluated the RanBP9, muskelin, and WDR26 domains required to mediate interactions with the CTLH complex. WDR26 and muskelin have a similar domain architecture, namely α-helical LisH, CTLH, and CRA motifs and a C-terminal ß-propellor domain – through WD40 repeats in WDR26 or kelch repeats in muskelin (**Supplemental Figure 2A**). Unique to muskelin, however, is a discoidin domain near the N-terminus. RanBP9 also has LisH, CTLH, and CRA motifs. We performed co-immunoprecipitation experiments with transfected RanBP9, muskelin, or WDR26 domain deletion constructs as baits (**Supplemental Figure 2B**). Though not required for MAEA association, the CTLH motif of RanBP9 was necessary for its interaction with both muskelin and WDR26 (**Figure 2A**). Deletion of the LisH motif of RanBP9, meanwhile, had no effect on muskelin and WDR26 association (**Figure 2A**).

**Figure 2.**
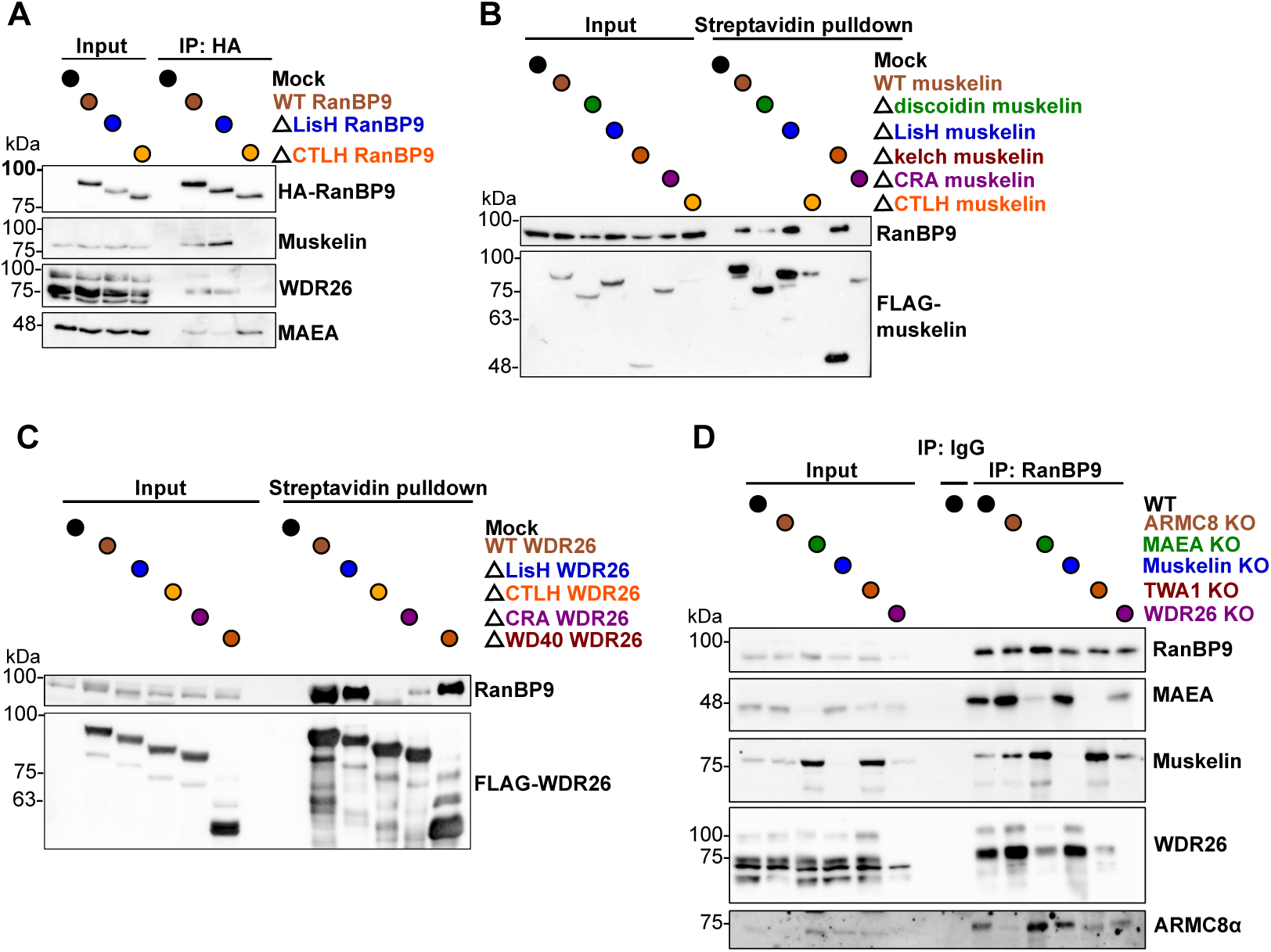
Muskelin and WDR26 associate with RanBP9 through the same surfaces. A) The indicated HA-RanBPM domain deletion constructs (see Supplemental Figure 2B) were transfected in shRanBPM HCT116 cells. After 24 hours, cells were lysed in whole-cell extract buffer and immunoprecipitated with an anti-HA antibody. Elutions were run on a western blot and analyzed for the presence of the indicated CTLH complex subunits. Experiments were performed in triplicate (n=3). B) The indicated FLAG-muskelin domain deletion constructs (containing a streptavidin binding peptide, see Supplemental Figure 2B) were transfected in muskelin KO HeLa cells. After 24 hours, cells were lysed in whole-cell extract buffer and muskelin complexes were isolated with streptavidin beads. Elutions were run on a western blot and analyzed for the presence of the indicated CTLH complex subunits. Experiments were performed in triplicate (n=3). C) The indicated FLAG-WDR26 domain deletion constructs (containing a streptavidin binding peptide, see Supplemental Figure 2B) were transfected in WDR26 KO HeLa cells. After 24 hours, cells were lysed in whole-cell extract buffer and WDR26 complexes were isolated with streptavidin beads. Elutions were run on a western blot and analyzed for the presence of the indicated CTLH complex subunits. Experiments were performed in triplicate (n=3). D) Immunoprecipitation of endogenous RanBPM. WT or indicated KO HeLa cells were lysed in whole-cell lysis buffer and subjected to immunoprecipitation with a RanBPM antibody that was crosslinked to beads. Elutions were run on a Western blot and analyzed for the presence of the indicated CTLH complex subunits. Experiments were performed in triplicate (n=3).

For muskelin, deletion of CTLH and CRA domains prevented binding to RanBP9 whereas deletion of the LisH, discoidin or kelch repeats did not (**Figure 2B**). Similarly, the CTLH and CRA domains of WDR26 were required for RanBP9 binding but the LisH and WD40 domains were not (**Figure 2C**). Next, we compared co-immunoprecipitations of endogenous RanBP9 across WT and KO cell lines of CTLH complex subunits. As expected, RanBP9 was still able to associate with CTLH complex subunits in either MKLN1 or WDR26 KO cells, and both muskelin and WDR26 still interacted with RanBP9 in any KO cell line other than their own (**Figure 2D**). These results are supported by similar co-IP experiments conducted for the *S. cerevisiae* complex as well as previously determined cryo-EM structures and *in vitro* binding assays for the human complex (Sherpa et al. 2021; van gen Hassend et al. 2023; Mohamed et al. 2021; Menssen et al. 2012). Overall, it is clear muskelin and WDR26 associate similarly with the CTLH complex via interactions between their CTLH-CRA domains with the CTLH-CRA domains of RanBP9.

### Muskelin can substitute for WDR26 as the ß-propellor link for supramolecular complex formation

It has yet to be determined in a cellular model whether WDR26 and muskelin co-exist as part of a supramolecular complex. Since the supramolecular complex contains two regions where WDR26 dimers or muskelin tetramers could bind to RanBP9, a “hybrid” supramolecular complex could in theory be formed with both moieties present together (**Figure 3A**). To address this possibility, transiently-transfected WT muskelin was immunoprecipitated from either DMSO or MG132-treated WT HeLa cells and co-immunoprecipitated WDR26 and RanBP9 levels were analyzed. Because of the muskelin autoregulation reported above, the co-immunoprecipitation was performed with lysates derived from MG132-treated cells to address the possibility that hybrid complexes are being degraded through the proteasome in nascent conditions. Although RanBP9 readily co-immunoprecipitated with overexpressed muskelin in both conditions, no detectable WDR26 was co-immunoprecipitated in either condition (**Figure 3B**). Similarly, a negligible amount of endogenous muskelin was co-immunoprecipitated with exogenous WDR26 in comparison to endogenous RanBP9 (**Figure 3C**), suggesting that the cell contains largely WDR26- or muskelin-specific supramolecular complexes. WDR26 is required *in vivo* for nuclear formation of the high order molecular complexes; however, we previously found that in WDR26 KO cells, proteasome inhibition by MG132 restored higher order nuclear complex formation (Onea et al. 2022). This indicated that if the CTLH subunits are not degraded in WDR26 KO cells, the supramolecular structure can still form even in the absence of WDR26.

**Figure 3.**
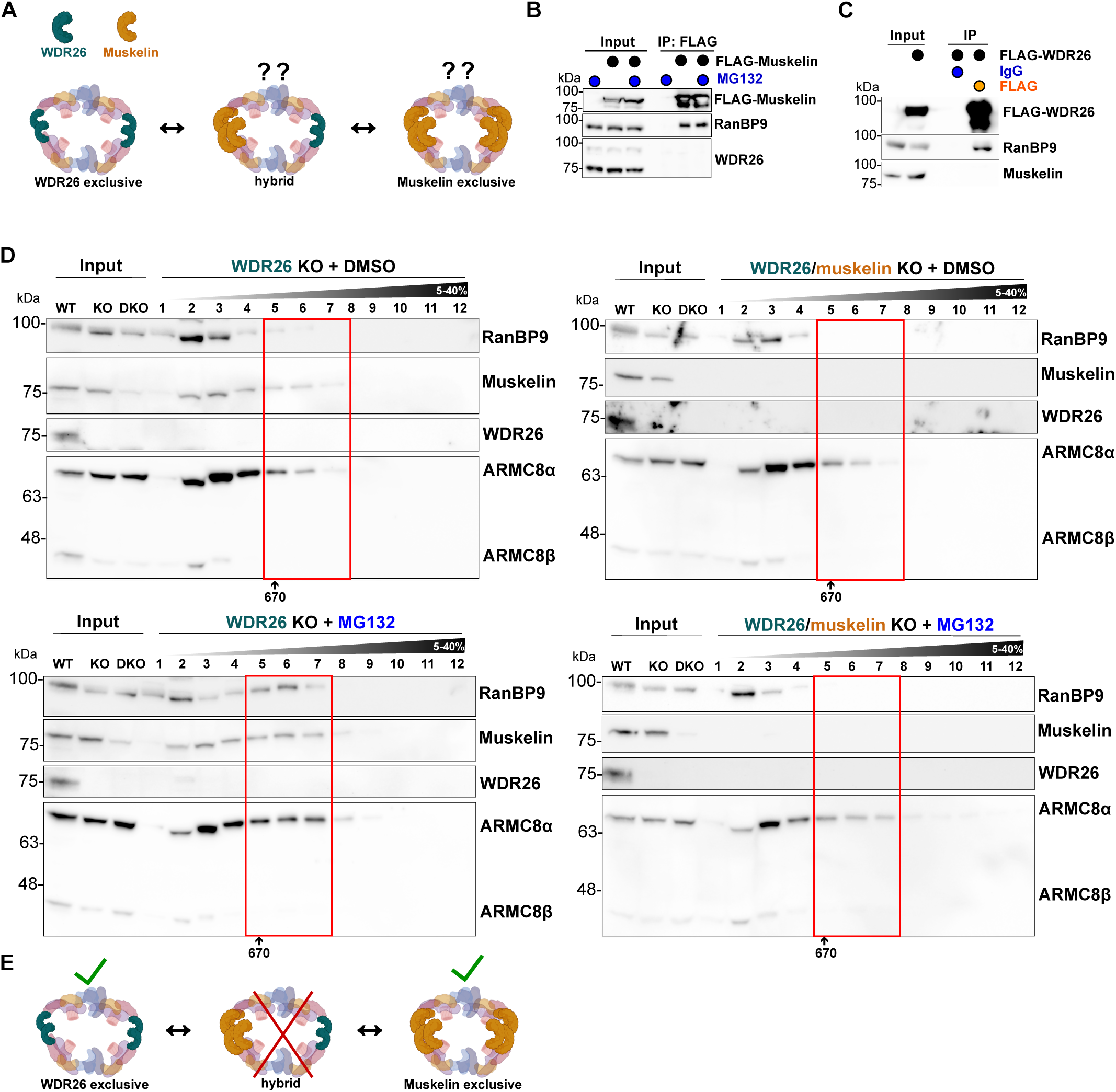
A muskelin exclusive supramolecular complex, but not a hybrid WDR26-muskelin complex, exists in cells. A) Schematic of possible muskelin or WDR26 combinations of CTLH supramolecular complexes. B) FLAG-muskelin or pcDNA3.1 empty vector was transfected in DMSO- or MG132-treated muskelin KO HeLa cells. After 24 hours, cells were lysed in whole-cell extract buffer and FLAG antibody was used for the co-immunoprecipitation assay. Elutions were run on a western blot and analyzed using the indicated antibodies. Experiments were performed in triplicate (n=3). C) FLAG-WDR26 or pcDNA3.1 empty vector was transfected in WDR26 KO HeLa cells. After 24 hours, cells were lysed in whole-cell extract buffer and FLAG or IgG antibody was used for the co-immunoprecipitation assay. Elutions were run on a western blot and analyzed using the indicated antibodies. Experiments were performed in triplicate (n=3). D) Whole-cell extracts were prepared from WDR26 KO and WDR26/muskelin double knockout (DKO) HeLa cells treated with DMSO or MG132 and separated by a 5–40% sucrose gradient. The resulting fractions were loaded on an SDS-PAGE gel, prepared for western blotting and analyzed with the indicated antibodies. Experiments were performed in triplicate (n=3). E) Updated schematic of CTLH supramolecular complex based on these results.

To test if muskelin is compensating for the lack of WDR26, we created a double muskelin/WDR26 KO cell line and performed sucrose gradients in the presence or absence of MG132. Sucrose gradients are comparable between WDR26 KO and muskelin/WDR26 KO cells in DMSO vehicle conditions, as has been observed previously (Sherpa et al. 2021); however, we found that while proteasome inhibition restored CTLH complex formation in WDR26 KO cells, it could not if both muskelin and WDR26 were absent (**Figure 3D**). We also tested whether muskelin or WDR26 were able to displace each other using sucrose gradients from WT cells transfected with either muskelin or WDR26. In untransfected cells, endogenous muskelin and WDR26 migrated mostly to fractions corresponding to supramolecular complex (fractions 5 and 6); however, upon transfection of exogenous muskelin, some of the endogenous WDR26 migrated to lower molecular mass fraction 3 and reciprocally, WDR26 transfection triggered some relocalization of muskelin into that same fraction (**Supplemental Figure 3A**). To summarize, muskelin and WDR26 can both serve as the ß-propellor link for the supramolecular CTLH complex, but a hybrid complex does not exist (**Figure 3E**). In addition, their association can be modulated by varying protein levels of either complex member (**Supplemental Figure 3B**).

### Muskelin instability requires both binding to RanBP9 and oligomerization

We next determined the requirements for muskelin to be regulated by the CTLH complex. We conducted CHX chase assays in WT and RMND5A KO cells transfected with 3 forms of muskelin. First, wild-type muskelin that can interact with RanBP9 and form a tetramer (WT), second, a mutant muskelin lacking the CRA domain that cannot interact with RanBP9 but can form a tetramer (ΔCRA), and finally a mutant lacking the LisH motif that can interact with RanBP9 but cannot form a tetramer (ΔLisH) (van gen Hassend et al. 2023; Delto et al. 2015). WT-muskelin was more unstable compared to either muskelin deletion mutant (**Figure 4**), demonstrating that muskelin that can form a supramolecular complex is degraded. This data supports the hypothesis that muskelin-containing supramolecular CTLH complexes are regulating complex-bound muskelin protein levels in an autoregulatory fashion.

**Figure 4.**
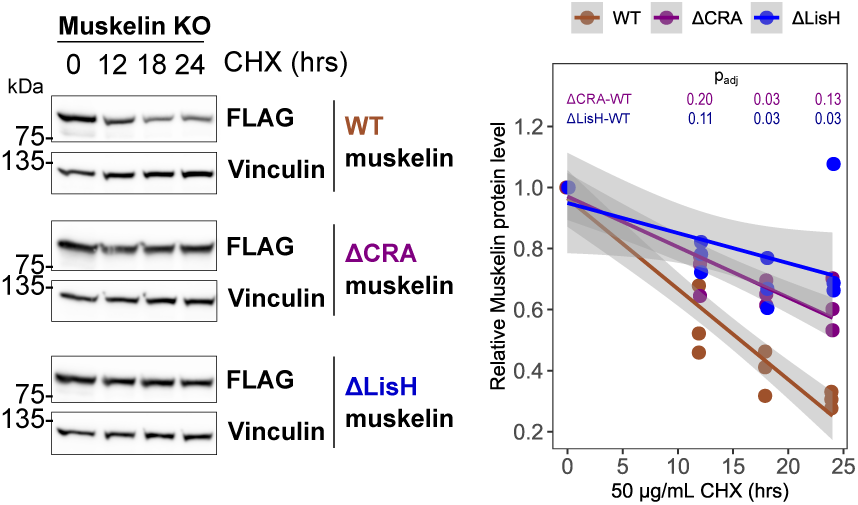
Muskelin instability depends on its LisH and CRA domains. Western blot quantification of muskelin protein levels in a cycloheximide (CHX) pulse-chase in muskelin KO HeLa cells. Representative western blots are shown (left) and quantification from n=3 independent experiments below. A Two-Way ANOVA revealed a significant interaction between 50 ug/uL CHX treatment time and muskelin constructs (F(2,28)=9.318, p=0.000791). Post-hoc comparisons using Tukey’s HSD test indicated that ΔCRA and ΔLisH significantly increased muskelin protein level at 18 and/or 24h CHX compared to WT (adjusted pvalue indicated on plot). Linear models were fit to the data using least squares method with each solid line indicating the best fit to the data and 95% confidence intervals indicated by the surrounding shaded area.

### Distinct proteome regulation by WDR26 and muskelin

Considering the observed dynamics between WDR26 and muskelin in binding to the CTLH complex, we thought that the differential association of these two subunits could result in CTLH complexes with distinct functions. Consequently, we performed a comprehensive proteomic analysis of their respective knockout cell lines to investigate potential functional differences. A principal component analysis of muskelin (n=5) and WDR26 (n=6) knockout proteomes revealed two discrete clusters delineated by their knockout status, wherein KO and control samples were segregated along the first principal component (PC1), accounting for approximately 70% of the observed variance in each case (**Supplemental Figure 4**). Overall, MKLN1 KO produced a less pronounced effect on global proteome regulation than WDR26 KO, since there were 132 significantly changed proteins out of 4218 in WDR26 KO, compared to only 39 changed proteins after loss of MKLN1 (adjusted p < 0.05, fold change cut-off of 1.5, **Figure 5A,B, Supplemental Data 3**). Since muskelin protein levels are decreased in WDR26 KO cells (Onea et al. 2022), it is not surprising that there were 16 significantly changed proteins in common between both proteomes, corresponding to 41% of all changed proteins in MKLN1 KO cells (**Figure 5C)**. Shared changed proteins included LIMA1 and ANPEP which both increased (**Figure 5D**), agreeing with their changes after either RMND5A or MAEA knockout (**Supplemental Figure 1B)**.

**Figure 5.**
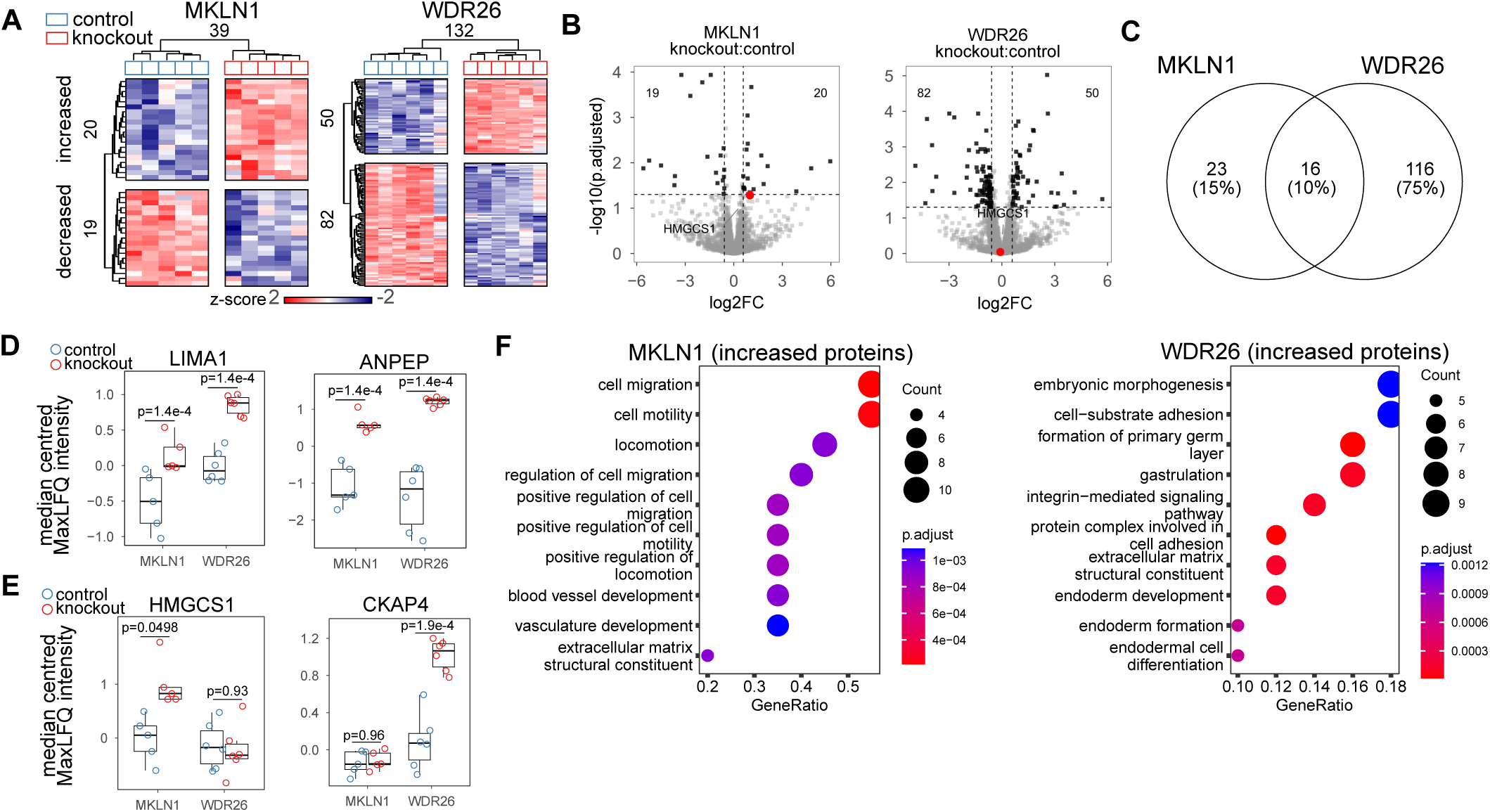
Distinct proteome regulation by muskelin and WDR26. A) Heatmap of differentially abundant proteins (adjusted p-value < 0.05, fold change > 1.5, statistics derived from limma, see methods) in whole-cell extracts of MKLN1 (n=5) or WDR26 KO (n=6) HeLa cells. Each row represents one protein and samples are represented in columns. Protein abundance is shown as z-scores quantified across rows. Hierarchical clustering of samples and proteins is shown. B) Volcano plot of all proteins with log2 fold change shown on the x-axis and −log10 adjusted p values shown on the y axis. MKLN1 KO vs respective controls on the left, WDR26 KO vs respective controls on the right. HMGCS1 is indicated in red. C) Overlap of proteins that had significant differential abundance in MKLN1 or WDR26 KO cells. D) Median centered intensity of proteins increased in both WDR26 and muskelin KO proteome samples, and in one (LIMA1) or both (ANPEP) of RMND5A and MAEA KO proteomes (see Supplemental Figure 1B). Boxplot midline indicates median values, bounds of the box indicate 25th and 75th percentiles, and maxima and minima indicate the largest point above or below 1.5 times the interquartile range. E) Median centered intensities for Pro/N-degron containing proteins, increased in RMND5A KO and/or MAEA KO (figure 1) proteomes that are increased in either MKLN1 (HMGCS1) or WDR26 (CKAP4) KO proteome samples. Boxplot midline indicates median values, bounds of the box indicate 25th and 75th percentiles, and maxima and minima indicate the largest point above or below 1.5 times the interquartile range. F) Top 10 GO terms functional annotation of proteins significantly increased in MKLN1 KO or WDR26 KO proteome samples. Gene ratio represents the fraction of genes in each GO category that were identified. P-values were adjusted using the Benjamini-Hochberg method.

In contrast to these similarities, many proteins showed specific regulation by either MKLN1 or WDR26. Two noteworthy examples are Pro/N-degron containing proteins CKAP4 and HMGCS1, which we previously found to be increased in RMND5A or MAEA knockouts (**Figure 5E**, **Figure 1E**). CKAP4 was not changed after MKLN1 knockout but increased in cells lacking WDR26 (**Figure 5E**). HMGCS1 levels, on the other hand, increased after MKLN1 knockout but were not changed in response to WDR26 knockout (**Figure 5B, E**).

To further investigate the specific functional differences in proteome regulation by muskelin and WDR26, we performed GO terms analysis on the list of increased proteins in each knockout. The top enriched GO terms associated with WDR26-increased proteins were associated with embryonic morphogenesis (GO:0048598). In contrast, the top ten significantly enriched GO terms (ranked by gene ratio) associated with MKLN1 increased proteins were largely associated with cell migration (GO:0016477) and cell motility (GO:0048870) (**Figure 5F**). Taken together, these findings are indicative of the distinct proteome regulation functions of MKLN1 and WDR26 acting in concert with the CTLH complex, mediated through mutually exclusive binding to the CTLH complex via RanBP9.

### The muskelin-containing supramolecular CTLH complex regulates HMGCS1 protein levels in an mTOR- and GID4-dependent manner

The proteome data determined that HMGCS1 was significantly increased in RMND5A KO cells (**Figure 1D**), and muskelin KO cells, but not WDR26 KO cells (**Figure 5B, E**). HMGCS1 contains a Pro/N-degron and we previously demonstrated it is regulated by CTLH complex substrate receptor GID4 (Owens et al. 2023). It therefore represents a potential CTLH target that may be distinctly regulated by muskelin and WDR26.

HMGCS1 was previously shown to be ubiquitinated and undergo rapid proteasomal degradation upon treatment with the mTOR inhibitor Torin-1, though an E3 ligase responsible for this was not identified (Zhao et al. 2015). By conducting a similar experiment with Torin-1 treatment in HeLa cells, we also observed rapid degradation of HMGCS1 (**Figure 6A)**. Remarkably, this degradation did not occur in RMND5A KO HeLa cells (**Figure 6A**), or in HeLa cells pre-treated with the GID4 chemical probe PFI-7 that occludes the GID4 binding pocket (Owens & Maitland et al., 2023) (**Figure 6B)**. As HMGCS1 was determined to be uniquely dependent on muskelin in the proteome datasets using unstimulated cells, we also compared the effect on HMGCS1 by Torin-1 treatment in WDR26 or muskelin KO HeLa cells. Expectedly, while Torin-1-induced HMGCS1 degradation in WDR26 KO cells was indistinguishable from WT cells, no degradation occurred in muskelin KO cells (**Figure 6A**). Together, this implicates the muskelin exclusive CTLH complex as the E3 responsible, via GID4 Pro/N-degron binding, for Torin-1 induced HMGCS1 ubiquitination and degradation.

**Figure 6.**
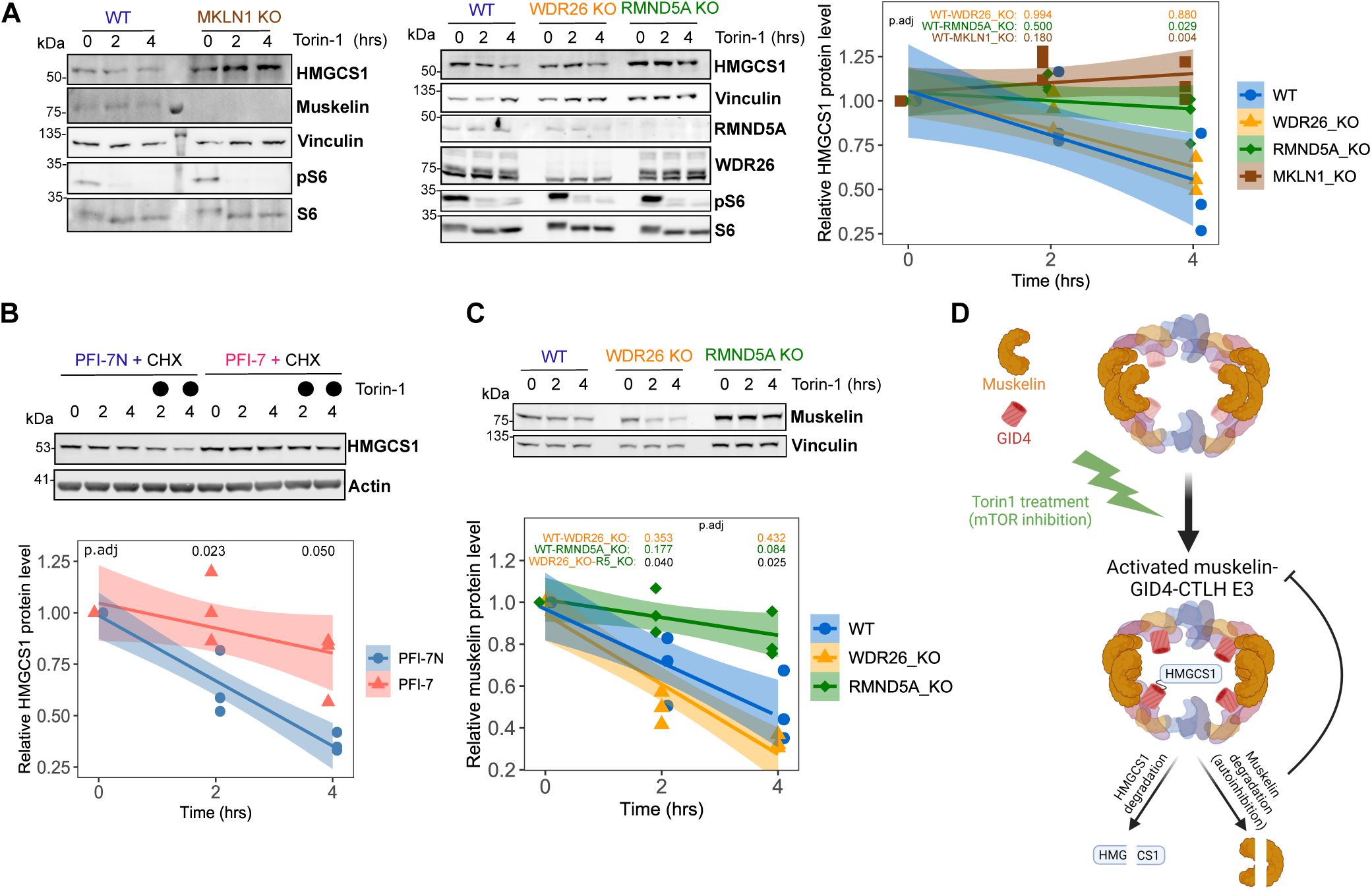
Torin-1 induced rapid degradation of HMGCS1 requires GID4 binding and the muskelin-CTLH complex. A) Whole-cell lysates were prepared from WT (blue, circle), MKLN1 KO (burgundy, square), WDR26KO (gold, triangle), or RMND5A KO (green, diamond) HeLa cells treated with 250 nM Torin-1 for the indicated time points and analyzed by western blot with the indicated antibodies. Quantification from n=3 independent experiments on the right. A Two-Way ANOVA revealed a significant interaction between treatment time and HMGCS1 levels (F(6,24)=3.902, p= 0.00734). Post-hoc comparisons using Tukey’s HSD test indicated that muskelin and RMND5A KO cells had significantly increased HMGCS1 protein level at 4 hr compared to WT (adjusted pvalues indicated on plot). Linear models were fit to the data using least squares method with each solid line indicating the best fit to the data and 95% confidence intervals indicated by the surrounding shaded area. B) Whole-cell lysates were prepared from WT HeLa cells pretreated for 24 hrs with 10 µM negative control compound PFI-7N (blue, circle) or PFI-7 (red, triangle). Cells were then treated with 50 µg/mL cycloheximide (CHX) or CHX and 250 nM Torin-1 for the indicated time points and analyzed by western blot with the indicated antibodies. Quantification from n=3 independent experiments on the right. A Two-Way ANOVA revealed a significant interaction between treatment time and HMGCS1 levels (F(2,12)=5.401, p= 0.021247). Post-hoc comparisons using Tukey’s HSD test indicated that PFI-7 treatment had significantly increased HMGCS1 protein level at 2 hr compared to PFI-7N (adjusted pvalues indicated on plot). Linear models were fit to the data using least squares method with each solid line indicating the best fit to the data and 95% confidence intervals indicated by the surrounding shaded area. C) The same lysates from (A) were run on western blot to assess muskelin levels. Quantification from n=3 independent experiments on the right. A Two-Way ANOVA revealed a significant interaction between treatment time and HMGCS1 levels (F(4,18)=6.302, p= 0.00236). Post-hoc comparisons using Tukey’s HSD test indicated that RMND5A KO cells had significantly increased HMGCS1 protein level at 2 and 4 hr compared to WDR26 KO (adjusted pvalues indicated on plot). Linear models were fit to the data using least squares method with each solid line indicating the best fit to the data and 95% confidence intervals indicated by the surrounding shaded area D) Schematic depicting the results presented here. The muskelin-CTLH complex is activated upon Torin-1 treatment and degrades HMGCS1 via GID4 binding. A negative feedback loop is present wherein muskelin is also degraded.

Curiously, Torin-1 treatment also caused muskelin levels to decrease in WT and WDR26 KO cells, but not in RMND5A KO cells (**Figure 6C**). These results were replicated in HEK293 cells, demonstrating that mTOR-mediated muskelin autoregulation is a general mechanism (**Supplemental Figure 5**). All together, these findings suggest that Torin-1 activates the muskelin-CTLH complex to degrade HMGCS1 using GID4 as the substrate receptor, but only briefly as muskelin autoregulation soon follows (**Figure 6D**).

## Discussion

The CTLH complex is a multi-subunit E3 ligase with recently reported chemical ligands to the substrate receptor GID4, generating interest for potential use of this E3 in targeted protein degradation (TPD) modalities (Owens et al. 2023; Chana et al. 2022; Yazdi et al. 2023; Schapira et al. 2019; Serebrenik et al. 2023). Prior to its implementation in TPD, however, the function and regulation of the CTLH complex must be better understood. Here, we identify a functional dichotomy between the ß-propeller subunits muskelin and WDR26 that translates into the formation of distinct CTLH complexes and correlates with the targeting of different substrates. Additionally, we provide the first evidence of a pathway that regulates the human CTLH complex activity. We demonstrate that the CTLH complex comprising muskelin, but not WDR26, rapidly degrades HMGCS1 upon mTOR inhibition, highlighting a biological regulation/pathway that affects the activity of a specific CTLH sub-complex. Muskelin degradation is also triggered, providing a mechanism for this complex to autoregulate its activity.

A recent report focused on GID4 in the presence or absence of its selective antagonist (PFI-7) identified HMGCS1 as the first human GID4 target (Owens et al. 2023). Here we show that HMGCS1 - a protein destabilized by the mTOR pathway (Zhao et al. 2015) - is degraded in a muskelin- and GID4-dependent manner when mTOR is inhibited. Since muskelin is also required for HMGCS1 degradation, its association with the core complex potentially creates the correct structure within the cavity of the supramolecular complex that facilitates GID4 anchoring of the HMGCS1 degron and efficient polyubiquitination of HMGCS1 by the CTLH catalytic subunits. This mechanism is likely similar to how Gid7 mediates Fbp1 degradation in yeast (Sherpa et al. 2021; Chrustowicz et al. 2023).

The *S. cerevisiae* Gid complex, homologous to the CTLH complex, achieves substrate diversity and regulation through distinct substrate receptors (Maitland et al. 2022). For example, the Gid complex can function through the substrate receptors Gid4, Gid10, and Gid11 that differ in their degron preferences, are activated by different stimuli, and undergo autoregulation once ubiquitination of targets is completed (Qiao et al. 2020; Menssen et al. 2018; Langlois et al. 2022; Kong et al. 2021).

Thus far, only GID4 substrate targeting has been defined for the human complex. Our data indicates that in order to regulate substrate diversity with only one substrate receptor, the CTLH complexes in multicellular organisms evolved to utilize and regulate the structurally similar but distinct β-propeller subunits WDR26 and muskelin. By way of their differing oligomeric states, the control of WDR26 versus muskelin complex formation dictates CTLH complex substrate preferences. This could be the reason why autoregulation of human CTLH complex activity occurs via its ability to degrade muskelin, allowing a quick shift between WDR26 and muskelin within the complex while maintaining availability of the general scaffold.

Regulated autoubiquitination is common amongst E3 ligases; for example, MDM2 autoubiquitinates itself when a threshold level of MDM2 protein is exceeded, preventing excess MDM2 and aberrant ubiquitination of non-substrate proteins (De Bie and Ciechanover 2011; Fang et al. 2000). Curiously, only muskelin – and not WDR26 – is regulated through autoubiquitination by the CTLH complex (Maitland et al. 2019; Jordan et al. 2023). One potential explanation may involve the structural differences between muskelin- and WDR26-containing CTLH complexes. For instance, muskelin has been reported to form a tetramer that can give rise to a sheet-like CTLH complex appearance while WDR26 dimerizes and mediates mainly donut-shaped supramolecular complexes (van gen Hassend et al. 2023). These structural differences potentially allow RING subunits MAEA and RMND5A to be in the proper position for muskelin ubiquitination, but not WDR26 (van gen Hassend et al. 2023; Sherpa et al. 2021). A recent study also demonstrated that the CTLH complex has intrinsic flexibility at the region where WDR26 and muskelin interact with RanBP9, potentially allowing muskelin to correctly orient itself for ubiquitination while the complex is formed (Chrustowicz et al. 2023). Although there is no evidence that WDR26 is regulated by CTLH complex-mediated ubiquitination activity, previous studies have shown that WDR26 protein levels correlate with changes in its gene expression during erythroid maturation (Zhen et al. 2020; Sherpa et al. 2022; An et al. 2014). This suggests that WDR26 protein levels may be regulated by specific signalling pathways at the level of gene expression.

Degradation of muskelin and muskelin-specific CTLH complex substrate HMGCS1 occurs simultaneously during mTOR inhibition. In this respect, autoubiquitination by the muskelin-containing CTLH complex appears mechanistically similar to what has been described for the Casitas B lineage (CBL) family of proteins. One member of this family, c-CBL, is an E3 ligase that regulates many processes including angiogenesis and EGFR signalling by targeting tyrosine kinases for degradation (Lyle et al. 2019). Phosphorylation of c-CBL mediated by its tyrosine kinase substrates changes the conformation of c-CBL, leading to simultaneous autoubiquitination and substrate ubiquitination (Ryan et al. 2006). A variety of post-translational modifications on muskelin, besides ubiquitination, have been detected in previous high-throughput screens, but as of now, the effects of these modifications on muskelin-containing CTLH complex activity or formation have not been explored (Hornbeck et al. 2015).

Our proteomic data in unstimulated HeLa cells highlighted HMGCS1 as a clear muskelin-specific target. We then demonstrated that HMGCS1 regulation by the CTLH complex was accelerated when mTOR was inhibited after finding evidence that mTOR regulates HMGCS1 protein levels (Zhao et al. 2015). In the future, other targets of such nature could be uncovered by conducting the proteomics in additional cell types and with a stimulus (e.g., using Torin-1 treatment).

Additionally, future studies could use our proteomics as a useful resource to characterize WDR26-containing CTLH complex targets. For example, one potential WDR26-specific target our proteomic data pointed towards was CKAP4. CKAP4 is a Pro/N-degron containing protein that was increased in RMND5A, MAEA, and WDR26 KO cells, but not muskelin KO cells. Investigating whether the WDR26-containing CTLH complex regulates CKAP4 protein levels, determining the cellular stimuli that may promote this regulation, and identifying the downstream effects of this regulation are intriguing topics for future studies.

Overall, we have integrated and extended the *in vitro* findings of yeast Gid7, its mammalian homologue WDR26, and the structurally similar muskelin in mediating supramolecular complex formation (Qiao et al. 2020; Mohamed et al. 2021; Sherpa et al. 2021; Maitland et al. 2019) into biological relevance using a human cell model. The biological importance of the interplay we uncovered between WDR26 and muskelin and how it affects CTLH complex function was evident in the case of Torin-1 induced degradation of HMGCS1. The increased activity of muskelin-containing CTLH complexes during mTOR inhibition provides clues into which biological processes require proper activation for normal function. The mTOR-AMPK axis regulates many processes including cellular proliferation, cilia development, glucose metabolism, and ribosome biogenesis, all of which have been proposed as CTLH complex-regulated processes in previous studies (Ling et al. 2020; Maitland et al. 2021; Hantel et al. 2022; Liu et al. 2020; Onea et al. 2022; McTavish et al. 2019). This interplay may also be relevant during human cytomegalovirus virus infection in which a WDR26-independent shift in complex composition was observed (Hashimoto et al. 2020) and for the drosophila CTLH complex in which the muskelin-dependent CTLH complex is responsible for degrading proteins as part of the maternal to zygotic transition (MZT), followed by muskelin ubiquitination and degradation (Cao et al. 2020; Zavortink et al. 2020). How this WDR26 and muskelin interplay and autoregulation may extend to other pathways and biological processes will be an exciting area of future study, and a necessary consideration for future development of CTLH complex-targeting TPD compounds.

## Materials and Methods

### Cell culture

HeLa and HEK293 cells were obtained from the American Type Culture Collection (ATCC). Generation of KO HeLa and HEK293 cell lines have been described previously (Onea et al. 2022; Maitland et al. 2019). All cells were cultured in high glucose (4.5g/L) Dulbecco’s modified Eagle’s medium (Wisent Bioproducts, St. Bruno, Quebec, Canada) supplemented with 10% fetal bovine serum, 1% sodium pyruvate, and 1% L-glutamine (609-065-EL, Wisent Bioproducts) at 37°C and 5% CO_2_. Cells were treated with DMSO (DMS666, Bioshop), 50 µM cycloheximide (Cyc003.1, Bioshop), 10 µM MG132 (474790, Cedarlane), 250 nM Torin1 (475991, Millipore Sigma), and 10 µM PFI-7N or PFI-7 (Structural Genomics Consortium) for the indicated times. Plasmid transfections were carried out with jetPRIME (CA89129-924, Polypus Transfection) on 80% confluent cells seeded the day before according to the manufacturer’s protocol. Cells were regularly tested to ensure absence of mycoplasma contamination.

### Cloning

All template DNA used in this study was described previously (Maitland et al. 2019), see Supplemental Table 1 below for a list of all cloning reagents. RanBP9 domain deletions have also been described previously (Salemi et al. 2015, 2014). Point mutations were generated via site directed mutagenesis with KOD polymerase (71086-3, Novagen) followed by Dpn1 digest (R0176, New England Biotechnology). Domain deletions were generated using tail-to-tail PCR with KOD polymerase and primers complementary to the regions immediately before and after the DNA to be deleted. All primers listed below were ordered from Integrated DNA Technologies (IDT). The entire cDNA sequence was validated by Sanger sequencing.

### Cell extracts

Whole cell extracts for immunoprecipitation and western blot were prepared by lysing cells in whole cell extract buffer containing fresh inhibitors (50 mM HEPES pH7.4, 150 mM NaCl, 10 mM EDTA, 10% glycerol, 0.5% NP40, 10 mM DTT, 1 mM Na3VO4, 10 mM NaF, 1 mM phenylmethylsulphonyl fluoride (PMSF), 1 μg/ml of aprotinin, 10 μg/ml of pepstatin, and 1 μg/ml of leupeptin) on ice for 20 minutes, then centrifuging for 20 minutes 13,000 rpm at 4°C, as previously described (Maitland et al. 2019). Whole cell extracts prepared for sucrose gradients underwent the same lysis but with lysis buffer containing no glycerol. For the PFI-7 experiment, cells were lysed in lysis buffer (20 mM Tris–HCl pH 8, 150 mM NaCl, 1 mM EDTA, 10 mM MgCl2, 0.5% Triton X-100, 12.5 U mL−1 benzonase). After 2 min incubation at RT, SDS was added to final 1% concentration. Preparation of nuclear extracts was also previously described (Onea et al. 2022); briefly, scraped cells were incubated in cytoplasmic lysis buffer (50 mM HEPES pH 7.4, 150 mM NaCl) supplemented with 10 µg/mL aprotinin, 2 µg/mL leupeptin, 2.5 µg/mL pepstatin, 1 mM DTT, 2 mM NaF, 2 mM Na3VO4, 0.1 mM PMSF, and 75 µg/mL digitonin (D141, Millipore Sigma). After a 10 minute incubation on ice, cells were centrifuged at 500xg for 1 minute and the supernatant was discarded. The resulting pellet was resuspended with nuclear lysis buffer (10 mM Tris-HCl pH 8.0, 100 mM NaCl, 0.1% sodium deoxycholate, 1 mM EDTA, 25% glycerol) supplemented with 10 µg/mL aprotinin, 2 µg/mL leupeptin, 2.5 µg/mL pepstatin, 1 mM DTT, 2 mM NaF, 2 mM Na3VO4, 0.1 mM PMSF, and 100 units/mL of Benzonase nuclease (E1014, Sigma Aldrich) and incubated on ice for 30 minutes. The samples were then centrifuged at 17000xg for 20 minutes and supernatants were isolated from cell debris. Protein concentration was estimated by Pierce 660 nm protein assay (22660, ThermoFisher).

### Immunoprecipitation

For RanBP9 IPs, preconjugation of antibody and immunoprecipitation protocol has been described elsewhere with slight modifications (Maitland et al. 2021). Four µg of RanBP9 antibody (F-1, sc-271727, Santa Cruz Biotechnology) and mouse IgG (sc-2025, Santa Cruz Biotechnology) were conjugated to 15 µL Dynabeads Protein G (10004D, Invitrogen) for 1 hour at 4°C with end-over-end rotation. Afterwards, antibody-bead conjugate was washed three times in 100 mM sodium Borate, pH 9 (M5162, Sigma-Aldrich), then resuspended in 20 mM DMP (Dimethyl pimelimidate dihydrochloride, D8388, Sigma-Aldrich) in 100 mM sodium Borate, pH 9 and rotated at room temperature for 30 minutes. Crosslinking was quenched by washing and incubation in 200 mM ethanolamine, pH 8 (S9640, Sigma-Aldrich) for 2 hours at 4°C with end-over-end rotation. After blocking, beads were washed three times in 1 mL whole cell extract buffer and kept on ice. 1 mg of whole cell extracts, prepared as described above, were then adjusted to 0.25% NP-40 and precleared using 5 µL Dynabeads Protein G. The preconjugated beads were then incubated with the precleared whole cell extracts overnight at 4°C while rotating. For FLAG and HA IPs, extracts were adjusted to 0.25% NP-40, pre-cleared and rotated overnight at 4°C with FLAG (M2, F1804, Sigma-Aldrich) or HA (HA-7, H9658, Sigma-Aldrich) then incubated for 1 hour at 4°C with Dynabeads protein G. In all IP types, beads were washed three times in wash IP buffer (25 mM HEPES pH 7.9, 60 mM KCl, 0.5 mM EDTA, 0.25% NP-40, 12% glycerol), resuspended in SDS loading buffer, and boiled at 95°C for 10 minutes to elute proteins prior to SDS-PAGE.

### Sucrose Gradient

Preparation of sucrose gradient fractions was described in a previous publication (Onea et al. 2022). Three hundred µg of either whole cell extract or nuclear extract, prepared as described above was loaded onto a 5-40% sucrose gradient (w/v) made with whole cell or nuclear lysis buffer the night before. Gradients were centrifuged in a SW41 rotor at 30 400 RPM for 16 hours at 4°C. Fractions were collected and 10% TCA and acetone were used to precipitate the proteins from each fraction. The protein pellets were resuspended in SDS loading buffer and boiled at 95°C for 10 min prior to SDS-PAGE.

### Western blot

For western blot using chemiluminescent imaging, protein was loaded onto a 10% or 8% SDS-polyacrylamide gel electrophoresis (SDS-PAGE) gel, transferred to Polyvinylidene difluoride (PVDF) membrane, hybridized, and assessed as described previously (Maitland et al. 2019). For fluorescence-based imaging, protein was loaded onto a 10% or 8% SDS-PAGE gel and was transferred to Immobilon-FL PVDF membrane (IPFL00005, Millipore). The blot was blocked for one hour and hybridized with primary antibody overnight in 0.5% fish gelatin (G7041, Sigma) dissolved in TBST (0.05% tween). The next day, the blot was washed 3 times in TBST for 10 minutes at room temperature and incubated in one of four secondary antibodies (IRDye 680RD Goat anti-Rabbit IgG, 926-68071, LI-COR Biotechnology; IRDye 680RD Donkey anti-Mouse IgG, 926-68072, LI-COR Biotechnology; IRDye 800CW Donkey anti-Rabbit IgG, 926-32213, LI-COR Biotechnology; IRDye 800CW Donkey anti-Mouse IgG, 926-32212, LI-COR Biotechnology) diluted at 1:10000 in 0.5% fish gelatin dissolved in TBST. After 3 washes in TBST for 10 minutes in the dark at room temperature, the blots were imaged using ChemiDoc MP (BioRad). Images were prepared and band intensities were quantified using ImageLab (BioRad). For immunoblotting of the PFI-7 experiment, samples were boiled in SDS loading buffer before western blotting using the NuPAGE electrophoresis and transfer system (Invitrogen). Immunoblots were imaged on a Li-Cor Odyssey CLx and quantified in Image Studio Lite v5.2.5 (Li-Cor Biosciences). Primary antibodies used in any experiments of this study are: Vinculin (E1E9V, Cell Signaling Technology); Muskelin (C-12, sc-398956, Santa Cruz Biotechnology); HA (HA-7, H3663 Sigma-Aldrich); FLAG (M2, F1804, Sigma-Aldrich); WDR26 (ab85962, Abcam); RanBP9 (5M, 71-001, Bio academia); ARMC8 (E-1, sc-365307, Santa Cruz Biotechnology); Tubulin (T5168, Sigma-Aldrich); RMND5A (custom, Yenzyme Antibodies); p-S6 (2215S, Cell Signaling Techonologies); S6 (2317S, Cell Signaling Technologies); Actin (A5441, Sigma-Aldrich); and HMGCS1 (A-6, sc-166763, Santa Cruz Biotechnology).

### MS Sample Preparation for Global Proteomics

HeLa cells at 75-80% confluency were trypsinized, cell pellet was collected by centrifugation, washed, and then frozen at −80°C. Cells were lysed by resuspension in 8 M urea, 50 mM ammonium bicarbonate (ABC), 10 mM Dithiothreitol (DTT), 2% Sodium dodecyl sulfate (SDS) and then sonicated with a probe sonicator (20×0.5 s pulses; Level 1). Twenty-five µg of protein lysate, as quantified by Pierce™ 660 nm Protein Assay (22660, 22663, ThermoFisher Scientific), was reduced in 10 mM DTT for 25 minutes, alkylated in 100 mM iodoacetamide for 25 minutes in the dark, followed by methanol precipitation as previously described (Kuljanin et al. 2017). The protein pellet was resuspended in 200 µL of 50mM ABC and subjected to a sequential digest first with 250 ng of LysC (125-05061, Wako Chemicals, USA) for 4 hours, then 500 ng of Trypsin/LysC (V5071, Promega) for 16 hours, followed by 500 ng of Trypsin (V5111, Promega) for an additional 4 hours. Digestions were incubated at 37°C at 600 rpm with interval mixing (30 seconds mix, 2 minutes pause) on a Thermomixer C (2231000667, Eppendorf). After the last digestion, samples were acidified with 10% formic acid (FA) to pH 3-4 and centrifuged at 14,000 x g to pellet insoluble material.

### diGLY enrichment

HeLa cells at 75-80% confluency were treated with 10 μM MG132 for 4 hours and processed exactly as described for global proteomics. After methanol precipitation, 1 mg protein was digested sequentially as described for global proteomics but with 6.66 μg of Lys-C, 20 μg of Trypsin/Lys-C, and 20 μg of Trypsin. Peptides were then dried using a SpeedVac (Thermo Scientific). PTMScan® Ubiquitin Remnant Motif (K-ε-GG) Antibody Bead Conjugate (5562, Cell Signaling Technology) (25 uL per sample) was crosslinked and used as previously described (Udeshi et al. 2013). Briefly, antibody-bead conjugate was washed three times in 100 mM sodium borate, pH 9 (M5162, Sigma-Aldrich), then resuspended in 20 mM DMP (Dimethyl pimelimidate dihydrochloride, D8388, Sigma-Aldrich) in 100 mM sodium borate, pH 9 and rotated at room temperature for 30 minutes. Crosslinking was quenched by washing and incubation in 200 mM ethanolamine, pH 8 (S9640, Sigma-Aldrich) for 2 hours at 4°C with end-over-end rotation. After blocking, beads were washed three times in 1 mL of 1x IAP buffer (provided by the kit) and kept on ice. Dried peptides were resuspended in 1 mL of IAP buffer, centrifuged, and the supernatant was added to the crosslinked antibody-beads and incubated with rotation for 1 hour at 4°C. After enrichment, beads were washed twice with IAP buffer, three times with PBS, and then peptides were eluted with two rounds of 5-minute incubations in 0.15% Trifluoroacetic acid (TFA). The eluted peptides were dried in a SpeedVac and reconstituted in 0.1% FA.

### Liquid Chromatography Tandem Mass-Spectrometry (LC-MS/MS) for Global Proteome and RMND5A diGly enrichment

Approximately 1 µg of peptide sample (as determined by Pierce BCA assay) was injected onto a Waters M-Class nanoAcquity HPLC system (Waters) coupled to an ESI Orbitrap mass spectrometer (Q Exactive plus, ThermoFisher Scientific) operating in positive mode. Buffer A consisted of mass spectrometry grade water with 0.1% FA and buffer B consisted of acetonitrile with 0.1% FA (ThermoFisher Scientific). All samples were trapped for 5 minutes at a flow rate of 5 µL/min using 99% buffer A and 1% buffer B on a Symmetry BEH C18 Trapping Column (5 mm, 180 mm x 20 mm, Waters). Peptides were separated using a Peptide BEH C18 Column (130 Å, 1.7 mm, 75 mm x 250 mm) operating at a flow rate of 300 nL/min at 35°C (Waters). Proteome Samples were separated using a non-linear gradient consisting of 1%–7% buffer B over 1 minute, 7%–23% buffer B over 179 minutes and 23%–35% buffer B over 60 minutes, before increasing to 98% buffer B and washing. RMND5A diGLY enriched samples were trapped for 5 minutes then separated using a non-linear gradient consisting of 1%–7.5% buffer B over 1 minutes, 7.5%–25% buffer B over 179 minutes, 25%–32.5% buffer B over 40 minutes and 32.5%-40% over 20 minutes before increasing to 98% buffer B and washing. MS acquisition settings are provided in Supplemental Table 2.

### Proteomic Data Analysis

The proteomic samples were processed using a label-free data-dependent acquisition (DDA) approach. Raw mass spectrometry files were converted to mzml format using msconvert within proteowizard (version 3.0.22167) (Chambers et al. 2012). Database searches were performed using MSFragger (version 3.7) (Teo et al. 2021; Kong et al. 2017) within FragPipe (version 19.0). For all database searches, MSFragger was used to search against the reviewed human proteome retrieved from Uniprot (2023-04-19 version) containing 40910 entries with reverse decoys and common contaminants added. Datasets were analysed in batches according to their acquisition time and analysis requirements. MKLN1 and WDR26 proteomes made up one analysis batch while the second batch consisted of RMND5A and MAEA proteomes. Ubiquitin remnant proteomic samples were processed in a separate run. Precursor and fragment mass tolerances were set to 20 ppm for total proteome. Enzyme specificity was set to trypsin, with up to two missed cleavages allowed for whole proteomes and three for ubiquitin remnant. Carbamidomethylation of cysteine (+57.02146 Da) was set as a fixed modification and oxidation of methionine (+15.9949 Da) and N-terminal acetylation (+42.0106 Da) were set as variable modifications. For ubiquitin remnant proteomics, GlyGly (+114.04293) was set as an additional variable modification. Filtering of peptide-spectrum matches (PSM) was performed using Philosopher (version 4.8.1) (da Veiga Leprevost et al. 2020) with false-discovery rate set to 1% at the PSM, peptide, and protein level. MSBooster was turned on (version 1.1.11) (Yang et al. 2023) and for ubiquitin remnant site localization, PTMProphet was used (Shteynberg et al. 2019) with a minimum site threshold of 0.5. Label-Free Quantification was performed by IonQuant (version 1.8.10) (Yu et al. 2021) using the MaxLFQ algorithm with match-between-runs enabled between all runs within each batch with a false-discovery rate threshold of 1%.

Subsequent data analysis was performed in R (version 4.1.2) with fully annotated analysis code available at https://github.com/d0minicO/CTLH_proteomes_analysis. DEP (version 1.16.0) (Zhang et al. 2018) was used for data normalization and imputation. Protein intensities for whole proteome data or peptide intensities for ubiquitin remnant data were filtered to retain rows missing in no more than two replicates of each condition. For whole proteome data, proteins with two or more peptides and at least one unique peptide was retained. MS1 intensities were log2-transformed and variance stabilizing normalization (vsn) was applied. Mixed imputation was performed to impute missing values with K-nearest neighbour (KNN) used for missing at random (MAR) and Quantile Regression Imputation of Left Censored Data (QRILC) used for missing not at random (MNAR). Limma (version 3.50.3) (Ritchie et al. 2015) was used to detect significant protein abundance changes with separate regression models for each batch. P-values were adjusted using Benjamini-Hochberg procedure with a significance threshold of 0.05 and fold-change threshold of 1.5. To visualise relevant biological variability across each dataset, dimensionality reduction was performed using principal components analysis (PCA) of a matrix containing all samples in each batch with all ubiquitin remnant-containing peptides or the subset of significantly changed proteins. Pathways analysis on sets of changed proteins was performed using clusterProfiler (version 4.2.2) (Wu et al. 2021) with all quantified proteins in each batch as background. Hierarchical clustering on heatmaps was done using the complete method within pheatmap (version 1.0.12) (https://github.com/raivokolde/pheatmap). To aid visualization of protein abundances in boxplots, median centering was applied.

### Statistics

Statistical analyses were performed using R (4.3.1). Statistical test used and associated metrics are defined in each figure legend. Results were considered significant when P < 0.05.

## Data Availability

The mass spectrometry proteomics data have been deposited to the ProteomeXchange Consortium via PRIDE (Perez-Riverol et al. 2019) partner repository.

## Acknowledgements

This work was supported by the Canadian Institutes of Health Research (MOP-142414 and PJT-169101 to C.S.P; FDN154328 to C.H.A); Mass spectrometry analyses were performed on equipment funded by a grant from the Canada Foundation for Innovation to G.A.L; G.O, M.E.R.M and B.G.C were supported by a Postgraduate Doctoral Scholarship from the Natural Sciences and Engineering Research Council of Canada. The Structural Genomics Consortium is a registered charity (no: 1097737) that receives funds from Bayer AG, Boehringer Ingelheim, Bristol Myers Squibb, Genentech, Genome Canada through Ontario Genomics Institute (OGI-196), EU/EFPIA/OICR/McGill/KTH/Diamond Innovative Medicines Initiative 2 Joint Undertaking (EUbOPEN grant 875510), Janssen, Merck KGaA (aka EMD in Canada and US), Pfizer and Takeda.

## Author Contributions

C.S.P, M.E.R.M. and G.O. designed project and experiments. M.E.R.M. and G.O. conducted most experiments, data analysis and generated most figures. B.C.G.C and X.W performed experiments. M.E.R.M. and D.D.G.O performed bioinformatic analyses and resulting figures. M.E.R.M. and G.O. wrote the manuscript with input from C.S.P. G.A.L, D.B.L, C.H.A. and C.S.P provided supervision and funding.

---

**Supplemental Table 1.**
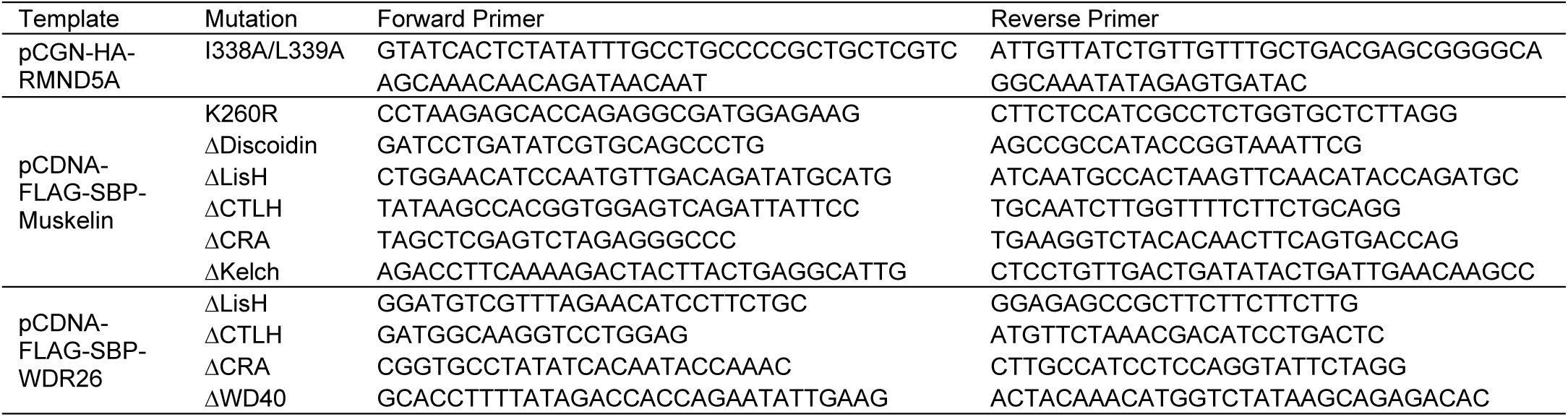
List of reagents used for mutagenesis.

**Supplemental Table 2.** Mass spectrometry scan settings.

| *Setting* | *Proteomes* | *RMND5A*<br>*diGLY*<br>*enrichment* |
| --- | --- | --- |
| *Orbitrap resolution (MS1)* | 70K | 70K |
| *Mass range* | 400-1500m/z | 400-1800m/z |
| *MS1 injection time* | 250ms | 250ms |
| *MS1 AGC target* | 3.00E+06 | 3.00E+06 |
| *Microscans* | 1 | 1 |
| *Data type* | Profile | Profile |
| *Lock mass* | 445.120025 | 445.12 |
| *MS2 resolution* | 17.5K | 17.5K |
| *Polarity* | Positive | Positive |
| *MS2 AGC target* | 2.00E+05 | 2.00E+05 |
| *MS2 injection time* | 64ms | 120ms |
| *Loop count* | 12 | 12 |
| *Isolation width* | 1.2m/z | 1.2m/z |
| *Isolation offset* | 0.5m/z | 0.5m/z |
| *MS2 Activation* | HCD | HCD |
| *Normalized Collision Energy* | 25 | 25 |
| *Dynamic exclusion* | enabled | enabled |
| *minimum AGC target* | 2.00E+03 | 2.00E+03 |
| *MS2 intensity threshold* | 3.10E+04 | 1.70E+04 |
| *Data dependent acquisition* | top12 | top12 |
| *Exclusion duration* | 30s | 30s |
| *Charge exclusion* | unassigned,<br>1, 7, 8, >8 | unassigned,<br>1, 7, 8, >8 |

**Supplemental Figure 1.**
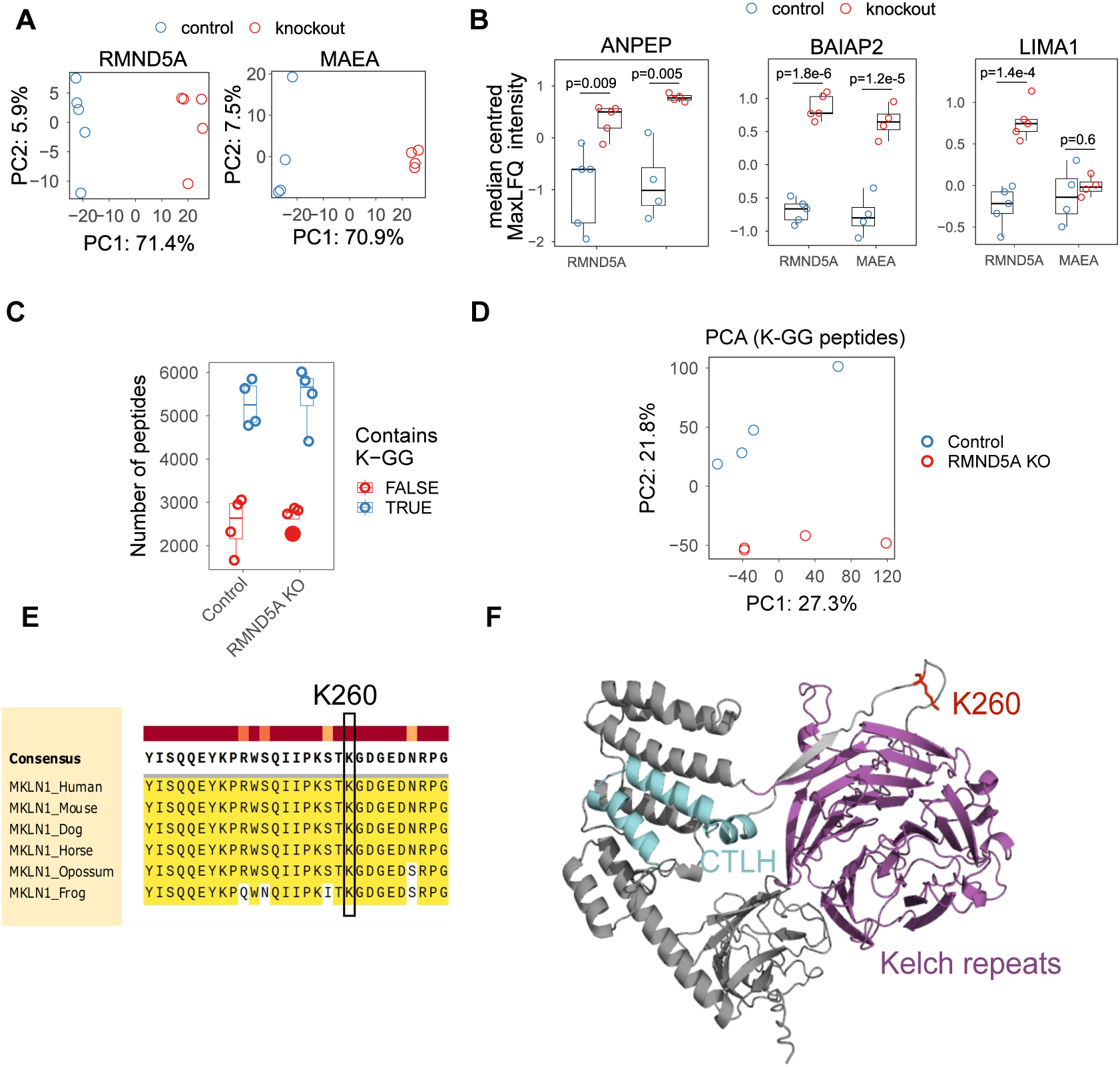
**Supplement to figure 1.** A) Principal component analysis of RMND5A KO (left) and MAEA KO (right) proteome samples with their respective control cell samples. B) Median centered intensities for specific proteins previously found to associate with the CTLH complex. Boxplot midline indicates median values, bounds of the box indicate 25th and 75th percentiles, and maxima and minima indicate the largest point above or below 1.5 * interquartile range. C) Relating to diGLY enriched proteome samples. Number of diGLY (K-GG)-containing peptides (blue) quantified in each sample versus not-diGLY containing peptides (red). D) Relating to diGLY enriched proteome samples. Principal component analysis of RMND5A KO (left) diGLY proteome samples with their respective control cell samples. E) Conservation of lysine 260 and surrounding residues of muskelin. F) Alphafold predicted structure of human muskelin (AF-Q9UL63-F1-model_v4) with CTLH motif in cyan, Kelch repeat in magenta, and residue K260 in red.

**Supplemental Figure 2.**
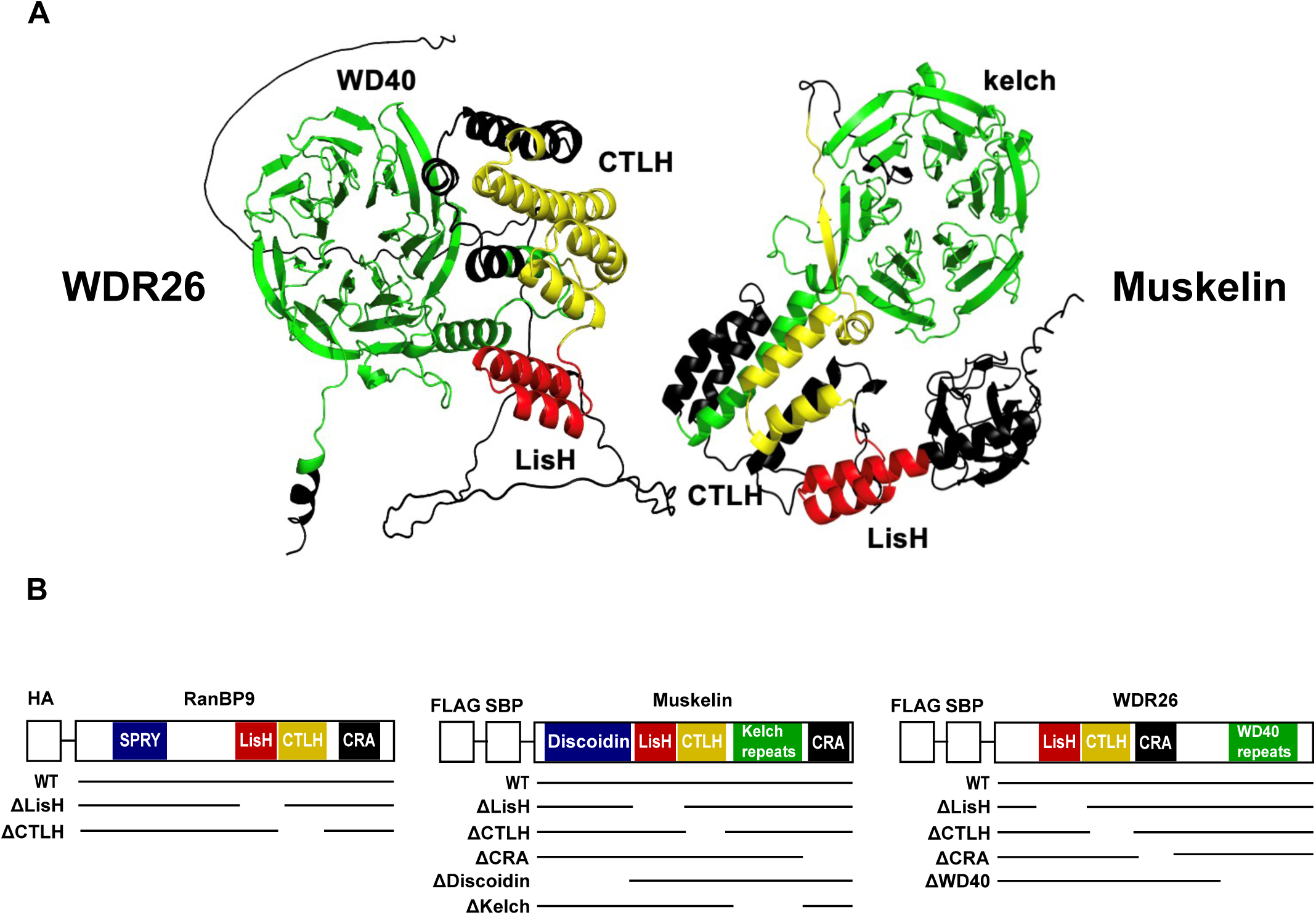
**Relating to figure 2.** A) Alphafold predicted structures of WDR26 or muskelin with their domain colour coded. LisH = red, CTLH = yellow, kelch or WD40 repeats = green. B) Organization of domain deletion constructs used in Figure 2A-C. SBP = streptavidin binding peptide.

**Supplemental Figure 3.**
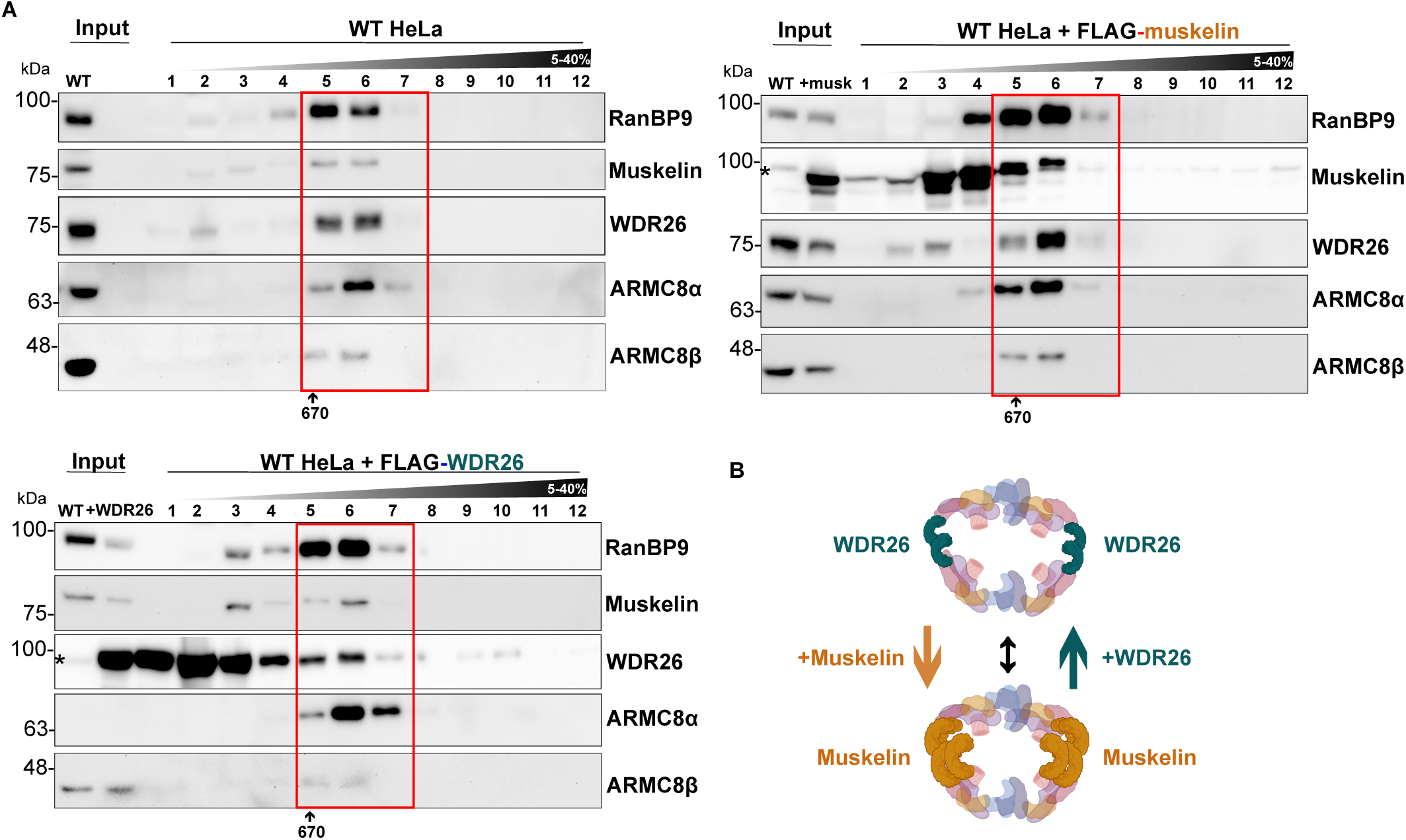
Overexpression of muskelin or WDR26 can displace the other β-propeller subunit in the CTLH supramolecular complex. A) Whole-cell extracts were prepared from WT HeLa cells transfected (24 hrs) with mock or the indicated FLAG tagged WDR26 or muskelin constructs and separated by a 5–40% sucrose gradient. The resulting fractions were loaded on an SDS-PAGE gel, prepared for western blotting and analyzed with the indicated antibodies. Experiments were performed in triplicate (n=3). B) Updated schematic of CTLH supramolecular complex demonstrating how the complex assembly can be influenced by overexpression of WDR26 or muskelin.

**Supplemental Figure 4.**
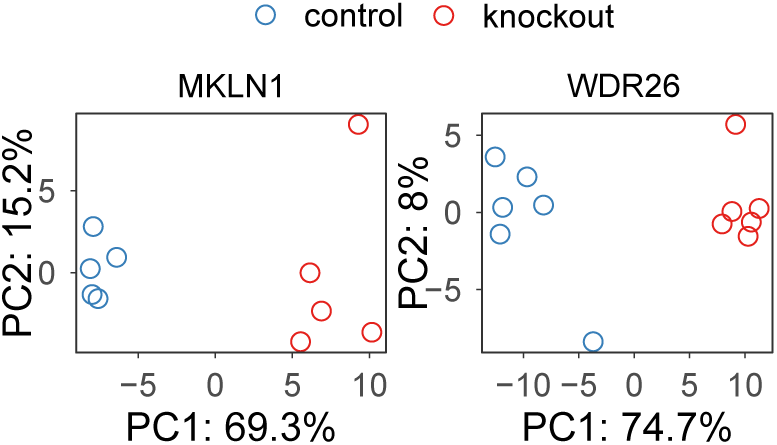
**Relating to Figure 5**. A) Principal component analysis of MKLN1 KO (left) and WDR26 KO (right) proteome samples with their respective control cell samples.

**Supplemental Figure 5.**
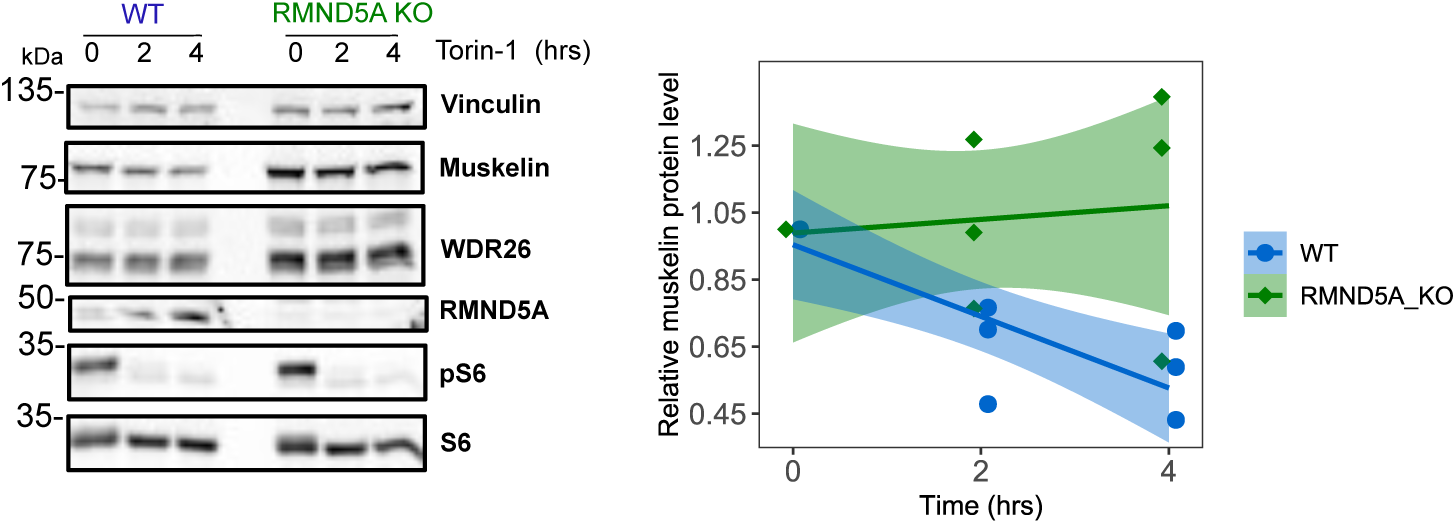
**Relating to figure 6.** **Torin-1 induced muskelin degradation in HEK293 cells.** A) Whole-cell lysates were prepared from WT (blue, circle) or RMND5A KO (green, diamond) HEK293 cells treated with 250 nM Torin-1 for the indicated time points and analyzed by western blot with the indicated antibodies. Quantification from n=3 independent experiments on the right. A Two-Way ANOVA did not reveal a significant interaction between treatment time and muskelin levels (F(2,12)=2.202, p= 0.1532). Linear models were fit to the data using least squares method with each solid line indicating the best fit to the data and 95% confidence i intervals indicated by the surrounding shaded area.

## References

An X, Schulz VP, Li J, Wu K, Liu J, Xue F, Hu J, Mohandas N, Gallagher PG. 2014. Global transcriptome analyses of human and murine terminal erythroid differentiation. Blood 123: 3466–3477.

Cao WX, Kabelitz S, Gupta M, Yeung E, Lin S, Rammelt C, Ihling C, Pekovic F, Low TCH, Siddiqui NU, et al. 2020. Precise Temporal Regulation of Post-transcriptional Repressors Is Required for an Orderly Drosophila Maternal-to-Zygotic Transition. Cell Rep 31: 107783.

Chambers MC, MacLean B, Burke R, Amodei D, Ruderman DL, Neumann S, Gatto L, Fischer B, Pratt B, Egertson J, et al. 2012. A cross-platform toolkit for mass spectrometry and proteomics. Nat Biotechnol 2012 3010 30: 918–920.

Chana CK, Maisonneuve P, Posternak G, Grinberg NGA, Poirson J, Ona SM, Ceccarelli DF, Mader P, St-Cyr DJ, Pau V, et al. 2022. Discovery and Structural Characterization of Small Molecule Binders of the Human CTLH E3 Ligase Subunit GID4. J Med Chem 65: 12725–12746.

Chen S-J, Wu X, Wadas B, Oh J-H, Varshavsky A. 2017. An N-end rule pathway that recognizes proline and destroys gluconeogenic enzymes. Science (80-) 355: eaal3655.

Chrustowicz J, Sherpa D, Li J, Langlois CR, Papadopoulou EC, Vu DT, Hehl LA, Karayel Ö, Beier V, von Gronau S, et al. 2023. Multisite phosphorylation dictates selective E2-E3 pairing as revealed by Ubc8/UBE2H-GID/CTLH assemblies. Mol Cell 1–16.

Chrustowicz J, Sherpa D, Teyra J, Loke MS, Popowicz GM, Basquin J, Sattler M, Prabu JR, Sidhu SS, Schulman BA. 2022. Multifaceted N-Degron Recognition and Ubiquitylation by GID/CTLH E3 Ligases. J Mol Biol 434: 167347.

da Veiga Leprevost F, Haynes SE, Avtonomov DM, Chang HY, Shanmugam AK, Mellacheruvu D, Kong AT, Nesvizhskii AI. 2020. Philosopher: a versatile toolkit for shotgun proteomics data analysis. Nat Methods 2020 179 17: 869–870.

De Bie P, Ciechanover A. 2011. Ubiquitination of E3 ligases: self-regulation of the ubiquitin system via proteolytic and non-proteolytic mechanisms. Cell Death Differ 2011 189 18: 1393–1402.

Delto CF, Heisler FF, Kuper J, Sander B, Kneussel M, Schindelin H. 2015. The LisH Motif of Muskelin Is Crucial for Oligomerization and Governs Intracellular Localization. Structure 23: 364–373.

Deshaies RJ, Joazeiro CAP. 2009. RING Domain E3 Ubiquitin Ligases. Annu Rev Biochem 78: 399–434.

Dikic I, Schulman BA. 2023. An expanded lexicon for the ubiquitin code. Nat Rev Mol Cell Biol 24: 273–287.

Dong C, Zhang H, Li L, Tempel W, Loppnau P, Min J. 2018. Molecular basis of GID4-mediated recognition of degrons for the Pro/N-end rule pathway. Nat Chem Biol 2018 145 14: 466–473.

Fang S, Jensen JP, Ludwig RL, Vousden KH, Weissman AM. 2000. Mdm2 is a RING finger-dependent ubiquitin protein ligase for itself and p53. J Biol Chem 275: 8945– 8951.

Hantel F, Liu H, Fechtner L, Neuhaus H, Ding J, Arlt D, Walentek P, Villavicencio-Lorini P, Gerhardt C, Hollemann T, et al. 2022. Cilia-localized GID/CTLH ubiquitin ligase complex regulates protein homeostasis of sonic hedgehog signaling components. J Cell Sci 135: 260203.

Hashimoto Y, Sheng X, Murray-Nerger LA, Cristea IM. 2020. Temporal dynamics of protein complex formation and dissociation during human cytomegalovirus infection. Nat Commun 2020 111 11: 1–20.

Hornbeck P V., Zhang B, Murray B, Kornhauser JM, Latham V, Skrzypek E. 2015. PhosphoSitePlus, 2014: mutations, PTMs and recalibrations. Nucleic Acids Res 43: D512–D520.

Huttlin EL, Bruckner RJ, Navarrete-Perea J, Cannon JR, Baltier K, Gebreab F, Gygi MP, Thornock A, Zarraga G, Tam S, et al. 2021. Dual proteome-scale networks reveal cell-specific remodeling of the human interactome. Cell 184: 3022–3040.e28.

Jordan VN, Ordureau A, An H. 2023. Identifying E3 Ligase Substrates With Quantitative Degradation Proteomics. ChemBioChem 24: 1–9.

Komander D, Rape M. 2012. The Ubiquitin Code. Annu Rev Biochem 81: 203–229.

Kong AT, Leprevost F V., Avtonomov DM, Mellacheruvu D, Nesvizhskii AI. 2017. MSFragger: ultrafast and comprehensive peptide identification in mass spectrometry–based proteomics. Nat Methods 2017 145 14: 513–520.

Kong K-YE, Fischer B, Meurer M, Kats I, Li Z, Rühle F, Barry JD, Kirrmaier D, Chevyreva V, San Luis B-J, et al. 2021. Timer-based proteomic profiling of the ubiquitin-proteasome system reveals a substrate receptor of the GID ubiquitin ligase. Mol Cell 81: 2460–2476.e11.

Kuljanin M, Dieters-Castator DZ, Hess DA, Postovit LM, Lajoie GA. 2017. Comparison of sample preparation techniques for large-scale proteomics. Proteomics 17.

Lampert F, Stafa D, Goga A, Soste MV, Gilberto S, Olieric N, Picotti P, Stoffel M, Peter M. 2018. The multi-subunit GID/CTLH E3 ubiquitin ligase promotes cell proliferation and targets the transcription factor Hbp1 for degradation. Elife 7.

Langlois CR, Beier V, Karayel O, Chrustowicz J, Sherpa D, Mann M, Schulman BA. 2022. A GID E3 ligase assembly ubiquitinates an Rsp5 E3 adaptor and regulates plasma membrane transporters. EMBO Rep 23: 1–17.

Ling NXY, Kaczmarek A, Hoque A, Davie E, Ngoei KRW, Morrison KR, Smiles WJ, Forte GM, Wang T, Lie S, et al. 2020. mTORC1 directly inhibits AMPK to promote cell proliferation under nutrient stress. Nat Metab 2020 21 2: 41–49.

Liu H, Ding J, Köhnlein K, Urban N, Ori A, Villavicencio-Lorini P, Walentek P, Klotz L-O, Hollemann T, Pfirrmann T. 2020. The GID ubiquitin ligase complex is a regulator of AMPK activity and organismal lifespan. Autophagy 16: 1618–1634.

Lyle CL, Belghasem M, Chitalia VC. 2019. c-Cbl: An Important Regulator and a Target in Angiogenesis and Tumorigenesis. Cells 2019, Vol 8, Page 498 8: 498.

Maitland MER, Kuljanin M, Wang X, Lajoie GA, Schild-Poulter C. 2021. Proteomic analysis of ubiquitination substrates reveals a CTLH E3 ligase complex-dependent regulation of glycolysis. FASEB J 35: e21825.

Maitland MER, Lajoie GA, Shaw GS, Schild-Poulter C. 2022. Structural and Functional Insights into GID/CTLH E3 Ligase Complexes. Int J Mol Sci 23: 5863.

Maitland MER, Onea G, Chiasson CA, Wang X, Ma J, Moor SE, Barber KR, Lajoie GA, Shaw GS, Schild-Poulter C. 2019. The mammalian CTLH complex is an E3 ubiquitin ligase that targets its subunit muskelin for degradation. Sci Rep 9: 1–14.

McTavish C, Bérubé-Janzen W, Wang X, Maitland M, Salemi L, Hess D, Schild-Poulter C. 2019. Regulation of c-Raf Stability through the CTLH Complex. Int J Mol Sci 20: 934.

Menssen R, Bui K, Wolf DH. 2018. Regulation of the Gid ubiquitin ligase recognition subunit Gid4. FEBS Lett 592: 3286–3294.

Menssen R, Schweiggert J, Schreiner J, Kušević D, Reuther J, Braun B, Wolf DH. 2012. Exploring the topology of the gid complex, the E3 ubiquitin ligase involved in catabolite-induced degradation of gluconeogenic enzymes. J Biol Chem 287: 25602– 25614.

Mohamed WI, Park SL, Rabl J, Leitner A, Boehringer D, Peter M. 2021. The human GID complex engages two independent modules for substrate recruitment. EMBO Rep 22: e52981.

Onea G, Maitland MER, Wang X, Lajoie GA, Schild-Poulter C. 2022. Distinct nuclear and cytoplasmic assemblies and interactomes of the mammalian CTLH E3 ligase complex. J Cell Sci 135.

Osugi Y, Fumoto K, Kikuchi A. 2019. CKAP4 Regulates Cell Migration via the Interaction with and Recycling of Integrin. Mol Cell Biol 39.

Owens DD., Maitland ME., Yazdi AK, Song X, Schwalm MP, Machado RA., Bauer N, Wang X, Szewczyk MM, Dong C, et al. 2023. A chemical probe to modulate human GID4 Pro/N-degron interactions. bioRxiv 2023.01.17.524225.

Perez-Riverol Y, Csordas A, Bai J, Bernal-Llinares M, Hewapathirana S, Kundu DJ, Inuganti A, Griss J, Mayer G, Eisenacher M, et al. 2019. The PRIDE database and related tools and resources in 2019: improving support for quantification data. Nucleic Acids Res 47: D442–D450.

Pfirrmann T, Villavicencio-Lorini P, Subudhi AK, Menssen R, Wolf DH, Hollemann T. 2015. RMND5 from Xenopus laevis Is an E3 Ubiquitin-Ligase and Functions in Early Embryonic Forebrain Development. PLoS One 10: e0120342.

Pickart CM. 2001. Mechanisms Underlying Ubiquitination. Annu Rev Biochem 70: 503– 533.

Popovic D, Vucic D, Dikic I. 2014. Ubiquitination in disease pathogenesis and treatment. Nat Med 20: 1242–1253.

Qiao S, Langlois CR, Chrustowicz J, Sherpa D, Karayel O, Hansen FM, Beier V, von Gronau S, Bollschweiler D, Schäfer T, et al. 2020. Interconversion between Anticipatory and Active GID E3 Ubiquitin Ligase Conformations via Metabolically Driven Substrate Receptor Assembly. Mol Cell 77: 150–163.e9.

Rape M. 2017. Ubiquitylation at the crossroads of development and disease. Nat Rev Mol Cell Biol 19: 59–70.

Ritchie ME, Phipson B, Wu D, Hu Y, Law CW, Shi W, Smyth GK. 2015. limma powers differential expression analyses for RNA-sequencing and microarray studies. Nucleic Acids Res 43: e47–e47.

Ryan PE, Davies GC, Nau MM, Lipkowitz S. 2006. Regulating the regulator: negative regulation of Cbl ubiquitin ligases. Trends Biochem Sci 31: 79–88.

Salemi LM, Almawi AW, Lefebvre KJ, Schild-Poulter C. 2014. Aggresome formation is regulated by RanBPM through an interaction with HDAC6. Biol Open 3: 418–430.

Salemi LM, Loureiro SO, Schild-Poulter C. 2015. Characterization of RanBPM Molecular Determinants that Control Its Subcellular Localization ed. A.F. Palazzo. PLoS One 10: e0117655.

Santt O, Pfirrmann T, Braun B, Juretschke J, Kimmig P, Scheel H, Hofmann K, Thumm M, Wolf DH. 2008. The Yeast GID Complex, a Novel Ubiquitin Ligase (E3) Involved in the Regulation of Carbohydrate Metabolism. Mol Biol Cell 19: 3323–3333.

Schapira M, Calabrese MF, Bullock AN, Crews CM. 2019. Targeted protein degradation: expanding the toolbox. Nat Rev Drug Discov 2019 1812 18: 949–963.

Serebrenik Y V., Mani D, Maujean T, Burslem GM, Shalem O. 2023. Pooled endogenous protein tagging and recruitment for scalable discovery of effectors for induced proximity therapeutics. bioRxiv 2023.07.13.548759.

Sherpa D, Chrustowicz J, Qiao S, Langlois CR, Hehl LA, Gottemukkala KV, Hansen FM, Karayel O, von Gronau S, Prabu JR, et al. 2021. GID E3 ligase supramolecular chelate assembly configures multipronged ubiquitin targeting of an oligomeric metabolic enzyme. Mol Cell 81: 2445–2459.e13.

Sherpa D, Mueller J, Karayel Ö, Xu P, Yao Y, Chrustowicz J, Gottemukkala K V, Baumann C, Gross A, Czarnecki O, et al. 2022. Modular UBE2H-CTLH E2-E3 complexes regulate erythroid maturation. Elife 11.

Shteynberg DD, Deutsch EW, Campbell DS, Hoopmann MR, Kusebauch U, Lee D, Mendoza L, Midha MK, Sun Z, Whetton AD, et al. 2019. PTMProphet: Fast and Accurate Mass Modification Localization for the Trans-Proteomic Pipeline. J Proteome Res 18: 4262–4272.

Sun X, Xie L, Qiu S, Li H, Zhou Y, Zhang H, Zhang Y, Zhang L, Xie T, Chen Y, et al. 2022. Elucidation of CKAP4-remodeled cell mechanics in driving metastasis of bladder cancer through aptamer-based target discovery. Proc Natl Acad Sci U S A 119.

Teo GC, Polasky DA, Yu F, Nesvizhskii AI. 2021. Fast Deisotoping Algorithm and Its Implementation in the MSFragger Search Engine. J Proteome Res 20: 498–505.

Udeshi ND, Mertins P, Svinkina T, Carr SA. 2013. Large-scale identification of ubiquitination sites by mass spectrometry. Nat Protoc 2013 810 8: 1950–1960.

van gen Hassend PM, Pottikkadavath A, Delto C, Kuhn M, Endres M, Schönemann L, Schindelin H. 2023. RanBP9 controls the oligomeric state of CTLH complex assemblies. J Biol Chem 299: 102869.

Vittal V, Stewart MD, Brzovic PS, Klevit RE. 2015. Regulating the Regulators: Recent Revelations in the Control of E3 Ubiquitin Ligases. J Biol Chem 290: 21244–21251.

Wu T, Hu E, Xu S, Chen M, Guo P, Dai Z, Feng T, Zhou L, Tang W, Zhan L, et al. 2021. clusterProfiler 4.0: A universal enrichment tool for interpreting omics data. Innovation 2: 100141.

Yang KL, Yu F, Teo GC, Li K, Demichev V, Ralser M, Nesvizhskii AI. 2023. MSBooster: improving peptide identification rates using deep learning-based features. Nat Commun 2023 141 14: 1–14.

Yazdi AK, Perveen S, Song X, Dong A, Szewczyk MM, Calabrese MF, Casimiro-Garcia A, Chakrapani S, Dowling MS, Ficici E, et al. 2023. Chemical Tools for the Gid4 Subunit of the Human E3 Ligase C-terminal to LisH (CTLH) Degradation Complex. bioRxiv 2023.11.13.566858.

Yu F, Haynes SE, Nesvizhskii AI. 2021. IonQuant Enables Accurate and Sensitive Label-Free Quantification With FDR-Controlled Match-Between-Runs. Mol Cell Proteomics 20: 100077.

Zavortink M, Rutt LN, Dzitoyeva S, Henriksen JC, Barrington C, Bilodeau DY, Wang M, Chen XXL, Rissland OS. 2020. The e2 marie kondo and the ctlh e3 ligase clear deposited rna binding proteins during the maternal-to-zygotic transition. Elife 9: 1– 48.

Zhang X, Smits AH, Van Tilburg GBA, Ovaa H, Huber W, Vermeulen M. 2018. Proteome-wide identification of ubiquitin interactions using UbIA-MS. Nat Protoc 2018 133 13: 530–550.

Zhao J, Zhai B, Gygi SP, Goldberg AL. 2015. MTOR inhibition activates overall protein degradation by the ubiquitin proteasome system as well as by autophagy. Proc Natl Acad Sci U S A 112: 15790–15797.

Zhen R, Moo C, Zhao Z, Chen M, Feng H, Zheng X, Zhang L, Shi J, Chen C. 2020. Wdr26 regulates nuclear condensation in developing erythroblasts. Blood 135: 208–219.

